# Biology and taxonomy of crAss-like bacteriophages, the most abundant virus in the human gut

**DOI:** 10.1101/295642

**Authors:** Emma Guerin, Andrey Shkoporov, Stephen R. Stockdale, Adam G. Clooney, Feargal J. Ryan, Thomas D. S. Sutton, Lorraine A. Draper, Enrique Gonzalez-Tortuero, R. Paul Ross, Colin Hill

**Affiliations:** aAPC Microbiome Ireland, University College Cork, Co. Cork, Ireland; bSchool of Microbiology, University College Cork, Co. Cork, Ireland; Teagasc Food Research Centre, Moorepark, Fermoy, Co. Cork, Ireland

**Keywords:** Bacteriophages, crAssphage, gut microbiota, human microbiome, phage taxonomy, phage characterisation

## Abstract

CrAssphage is yet to be cultured even though it represents the most abundant virus in the gut microbiota of humans. Recently, sequence based classification was performed on distantly related crAss-like phages from multiple environments, leading to the proposal of a familial level taxonomic group [Yutin N, et al. (2018) Discovery of an expansive bacteriophage family that includes the most abundant viruses from the human gut. Nat Microbiol 3(1):38–46]. Here, we assembled the metagenomic sequencing reads from 702 human faecal virome/phageome samples and obtained 98 complete circular crAss-like phage genomes and 145 contigs ≥70kb. *In silico* comparative genomics and taxonomic analysis was performed, resulting in a classification scheme of crAss-like phages from human faecal microbiomes into 4 candidate subfamilies composed of 10 candidate genera. Moreover, laboratory analysis was performed on faecal samples from an individual harbouring 7 distinct crAss-like phages. We achieved propagation of crAss-like phages in *ex vivo* human faecal fermentations and visualised *Podoviridae* virions by electron microscopy. Furthermore, detection of a crAss-like phage capsid protein could be linked to metagenomic sequencing data confirming crAss-like phage structural annotations.

**Significance:** CrAssphage is the most abundant biological entity in the human gut, but it remains uncultured in the laboratory and its host(s) is unknown. CrAssphage was not identified in metagenomic studies for many years as its sequence is so different from anything present in databases. To this day, it can only be detected from sequences assembled from metagenomics or viromic datasets (**crAss** – **cr**oss **Ass**embly). In this study, we identified 243 new crAss-like phages from human faecal metagenomic studies. Taxonomic analysis of these crAss-like phages highlighted their extensive diversity within the human microbiome. We also present the first propagation of crAssphage in faecal fermentations and provide the first electron micrographs of this extraordinary bacteriophage.

## Introduction

In recent years, increasing numbers of bacteria, archaea, fungi, protists and viruses residing on and within the human body have been associated with various states of human health and disease, including diet, age, weight, inflammatory bowel disease (IBD), diabetes, and cognition (1–7). A relatively small number of eukaryote viruses present in the gastrointestinal tract can target the human host, however, much larger and much more complex populations of viruses that target bacteria (bacteriophages) also reside there. The role of phages in the gut has been a subject of increased interest as initial investigations have revealed substantial differences in bacteriophage populations between healthy and diseased cohorts (7–11). It is likely that phages have an important role in shaping our gut microbiome, but their precise role remains poorly understood.

In 2014, metagenomic studies of the viral fraction of the human gut microbiota identified a DNA phage, crAssphage, detectable in approximately 50% of individuals from specific human populations and reaching up to 90% of the total viral DNA load in faeces of certain individuals (12). Dutilh and colleagues noted that crAssphage had been overlooked in previous metagenomic studies as the vast majority of its genes do not match known sequences present in databases. It has been predicted, based on indirect evidence using host co-occurrence profiling, that prototypical crAssphage infects *Bacteroides*, an abundant genus of bacteria important for the normal gut function of humans. However, since crAssphage has never been isolated in culture, its host range, replication strategy, virion morphology and impact on the human gastrointestinal microbiota remains unknown. Thus, a better understanding of crAssphage is crucial to understanding phage host dynamics in the human gut microbiota.

Originally crAssphage was published as an individual phage following cross-assembly of several metagenomic samples (12). Analysis by Manrique *et al*., of the healthy human gut phageome identified 4 circular crAssphage genomes and several related incomplete contigs (10). PCR amplification and sequencing of the crAssphage polymerase gene by Liang and colleagues similarly demonstrated diversity amongst crAssphage-positive faecal samples (13). Recently, Cinek *et al*. described updated PCR primer sequences for the detection and evaluation of crAssphage diversity, while Stachler *et al*. developed their own primers targeting conserved genomic regions to evaluate the abundance of crAssphage as an indicator of human faecal pollution (14, 15). Finally, an epidemiological survey of crAssphages conducted by Dutilh, Edwards and colleagues has suggested crAssphage is associated with humans and primates globally with significant diversity (manuscript currently in preparation).

A recent study provided the first detailed sequence-based taxonomic categorisation of crAss-like phages, proposing a novel familial level taxonomic group that would include crAssphage itself, as well as various related bacteriophages, from multiple environments (16). However, the authors noted that this classification is in contrast with the classical viral taxonomy scheme currently in use. Such taxonomy strictly categorises crAssphage as a member of the *Podoviridae* family. Previous attempts to reconcile sequence-based and classical viral taxonomy have proposed *Podoviridae* sharing >40% orthologous protein-coding genes be grouped at the taxonomic rank of genus, while phages sharing only 20-40% orthologous protein-coding genes should be grouped at the higher taxonomic rank of subfamily (17). Other reports describe a phage genus as a cohesive group of viruses sharing >50% nucleotide sequence similarity (18). As crAssphage is not a single entity, but rather a group of crAss-like phages that share similarity with the prototypical crAssphage at various levels, a comparative analysis of crass-like phage sequences is required to enable detailed taxonomic characterisation.

In this study, we combine several *in silico* and *in vitro* approaches to further explore the diversity of crAss-like phages in the human gut, and better understand their biological properties. We performed an in-depth analysis of crAss-like sequences from a number of previously published and unpublished human faecal virome datasets (1, 7, 9, 10, 19). Subsequent to the assembly of metagenomic sequencing reads, crAss-like phage contigs were identified using conserved genetic signatures. In total, 98 complete circular and 145 near-complete (≥70kb) linear contigs of crAss-like phages were identified for genomic and taxonomic analyses. Laboratory analysis of crAss-like phages was focused on a human donor identified as a stable carrier of several highly predominant crAssphage-like DNA sequences, including one closely related to the prototypical crAssphage. *Ex vivo* faecal fermentations enabled the amplification of a virus highly related to the prototypical crAssphage, with electron micrographs supporting the proposal that crAss-like phages are members of the *Podoviridae* family. These results represent the first example of biological characterisation of this highly prevalent and, potentially, very important human microbiome virus.

## Results

### Detection of crAss-like phage contigs

Following the assembly of 702 human faecal virome/phageome metagenomic samples listed in Supplementary Table 1, contigs were screened for relatedness to the prototypical crAssphage virus, henceforth referred to as crAssphage *sensu stricto*. Initially, the polymerase of crAssphage *sensu stricto* (UGP_018, NC_024711.1) was used for crAss-like phage detection due to its use in several studies as a genetic signature to determine diversity of crAss-like phages (13, 20, 21). However, we extended our criteria in order to include partial genomes (≥70kb) that may not have included the polymerase gene in the assembly. Therefore, after an initial detection of crAss-like phages using the polymerase sequence, we identified the most conserved crAss-like phage protein in our dataset as the terminase protein, encoded by crAssphage *sensu stricto* UGP_092. The terminase was subsequently used as a second genetic signature for identifying crAss-like phage contigs.

Initially, 239 contigs ≥70kb were detected with similarity to crAssphage *sensu stricto* polymerase sequence. An additional 59 contigs ≥70kb were subsequently detected with relatedness to crAssphage *sensu stricto* terminase sequence. Following an initial examination of the contig sequences retrieved, more stringent parameters were implemented. Only contigs whose polymerase and/or terminase sequence(s) aligned with greater than 350bp were considered for further analysis as crAss-like phages. This reduced the total number of crAss-like phages to 256. In addition, as several assembled metagenomic samples were from the same person sequenced at multiple time points, redundant contigs were removed from further analysis. When two or more contigs aligned with 100 percent identity, the longer contig or the contig with the highest coverage was retained. This resulted in a total of 244 crAss-like contigs (including crAssphage *sensu stricto*), with 143 contigs containing both a polymerase and terminase, 60 a polymerase only and 40 a terminase only. Of the 244 crAss-like phage contigs, metadata was available for the majority of their originating faecal samples. CrAss-like phages were detected in healthy individuals across a wide age range (including infants 1 year of age and individuals ≥65 years of age) and individuals suffering from Crohn’s disease, ulcerative colitis, cystic fibrosis, kwashiorkor and marasmus.

### Taxonomy of crAss-like phages

In order to compare the phylogeny of the more distantly related phages proposed to be included into a crAss-like familial level taxon by Yutin *et al*. (16) with those identified in this study, a phylogenetic tree of conserved crAss-like phage terminase sequences was constructed (Supplementary Figure 1). Amino acid terminase sequences were used to generate mid-point rooted phylogenetic trees. Predominantly, the terminase sequences of very distant crAss-like phage relatives identified by Yutin *et al*. from various environmental sources were distinct from the various candidate genera of crAss-like phages observed in the phylogram. However, the human gut microbiome phage, IAS virus (16), characterised by Yutin *et al*. as crAss-like, clustered closely with candidate genus VI crAss-like phages identified in this study.

**Figure 1.**
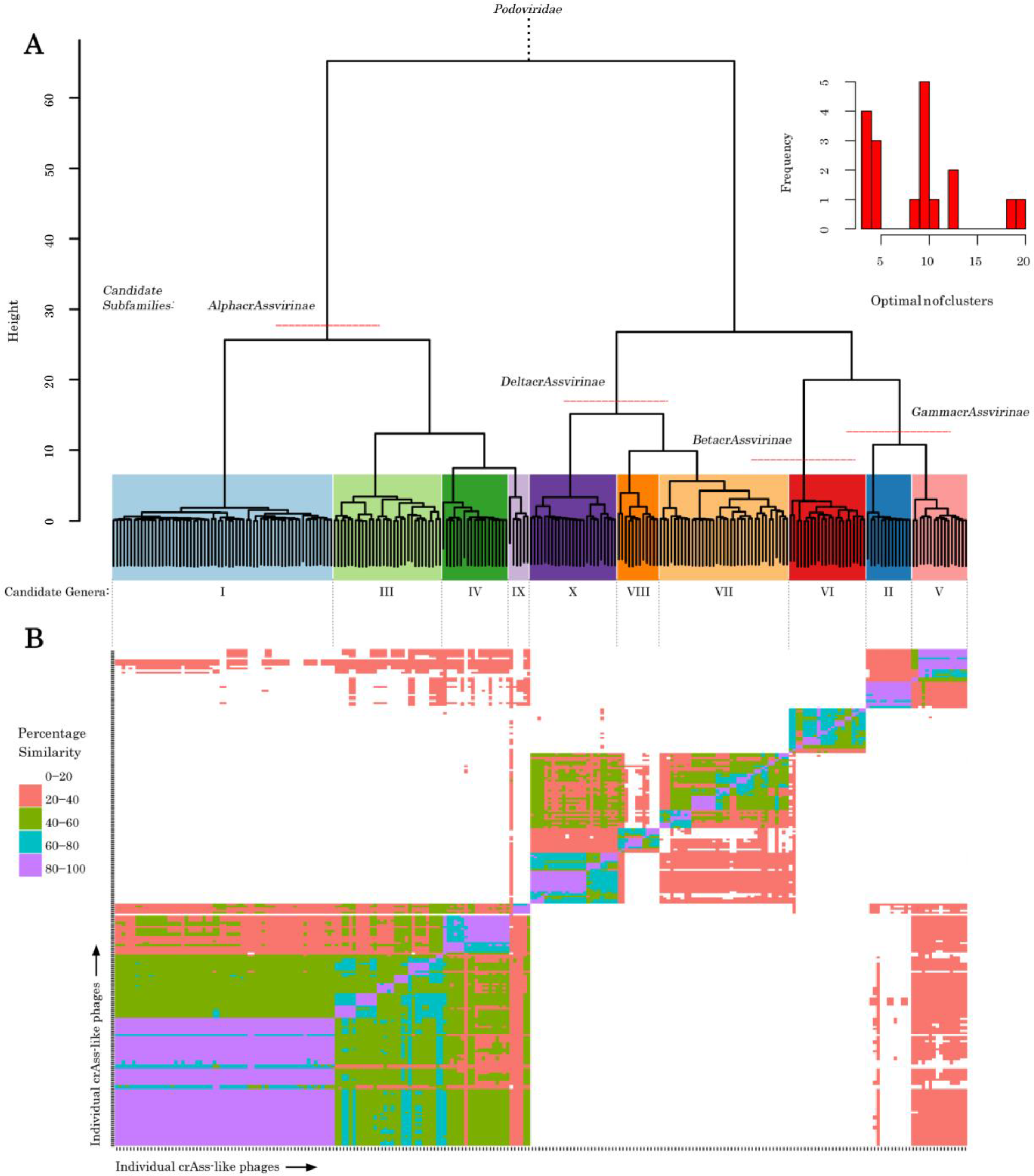
Determination of crAssphage candidate subfamilies and genera based on the percentage of shared protein-encoding genes. **(A)** The 4 red lines cut the hierarchical clustering dendrogram of crAss-like phage contigs, with Euclidean distances calculated between the percentages of shared protein-encoding genes, into the 4 proposed candidate subfamilies of crAss-like phages. The histogram insert (top-right) represents the calculated optimal number of crAss-like phage clusters. The 10 optimal crAss-like phage clusters represent the putative candidate genera, and are assigned specific colours. **(B)** Heatmap showing the percentage of shared protein-coding genes between crAss-like phage genomes. CrAss-like phages with 20-40% shared protein encoding genes are considered related at the subfamily level while phages with >40% similarity are believed to be related at the genus level, consistent with the calculated number of crAss-like phage clusters.

Previously, studies have used the percentage of shared homologous proteins as a means of defining phage taxonomic ranks (17). Therefore, clusters of phages sharing between 20-40% of their protein-coding genes were categorised as related at the subfamily level, while phages sharing >40% protein-coding genes were grouped at the genus level. A heatmap based on the percentages of shared orthologous proteins suggests that crAss-like phages form 4 candidate subfamilies. The four subfamilies were assigned the nomenclature *alphacrAssvirinae* (which contains crAssphage *sensu stricto*), *betacrAssvirinae* (which contains IAS virus), *gammacrAssvirinae* and *deltacrAssvirinae* (Figure 1). These subfamilies can be further subdivided into 10 candidate genera, with Candidate Genus I containing crAssphage *sensu stricto* and Candidate Genus VI containing the IAS virus. Metadata of all crAss-like phages analysed in this study, including their categorisation into the various taxonomic divisions, is available in Supplementary Table 2.

An alternative approach for characterising the encoded proteome of crAss-like phages was performed by visualisation of genome clusters using the t-SNE machine learning algorithm with Euclidean distances of orthologous genes distribution between genomes as an input. Applying the previously determined 10 crAss-like phage candidate genera classifications to the t-SNE two-dimensional ordination demonstrated that some clusters showed uniformity while others groups were quite dispersed, such as Candidate Genus II and VII, respectively (Figure 2A). In addition, no single cluster of crAss-like phages is exclusively associated with healthy or diseased individuals.

**Figure 2.**
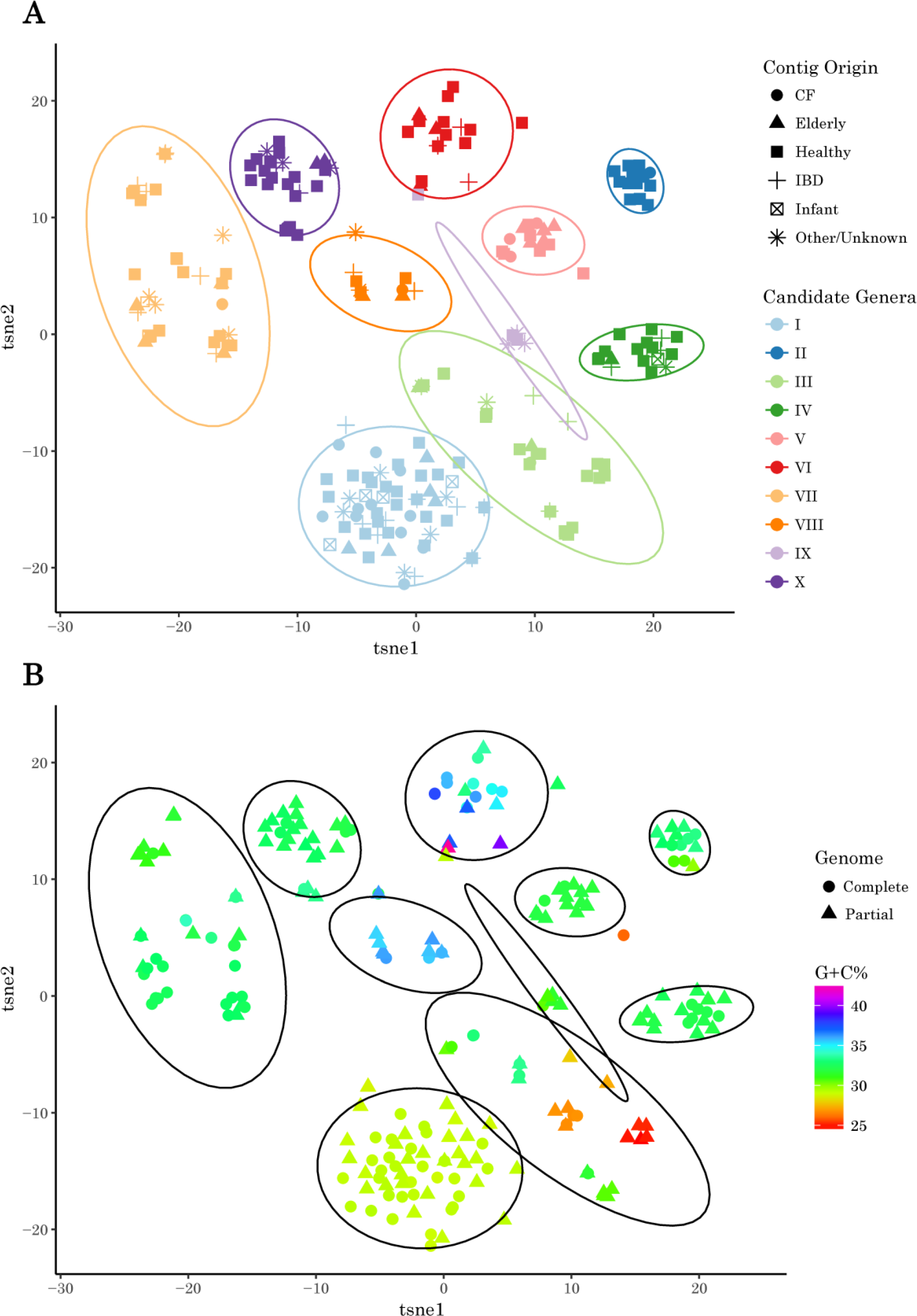
Two-dimensional ordination of crAss-like phages based on the abundance of their protein-encoded orthologous sequences was performed using t-SNE machine learning algorithm. **(A)** CrAss-like phages are coloured by candidate genus annotations and shape is determined by their origin. CrAss-like phages originating from individuals with kwashiorkor and marasmus, or lacking metadata, are grouped together as ‘Other/Unknown’. **(B)** CrAss-like phages are coloured by the percentage G+C nucleotide composition of their contig, while shape represents complete (circular) or partial (linear) genomes.

Groups of crAss-like phages with a similar G+C nucleotide content would be expected to infect related bacteria, since phage G+C content often aligns to that of its host (22, 23). Therefore, several groups of crAss-like phages, such as candidate genera II, IV, V, VII and X, are likely infect closely related bacterial taxa within the human microbiome (Figure 2B). Candidate genus I is the most homogenous group of crAss-like phages containing crAssphage *sensu stricto* and 30 additional complete circular genomes and 29 linear contigs ≥70kb with a distinct G+C nucleotide content (29.11 ± 0.14%). Candidate genera III and VI display the greatest heterogeneity, with G+C contents of 28.94 ±3.03% and 35.81 ± 2.56%, respectively.

### Nucleotide comparison of crAss-like phages

To further investigate the relatedness of crAss-like phages, a more detailed comparison at the nucleotide level was performed by calculating their average nucleotide identity (Figure 3). Candidate genera III and VI of crAss-like phages, as defined by the percentage of their shared encoded proteins, also do not cluster into clearly definable groups based on nucleotide composition. Candidate Genus I, containing crAssphage *sensu stricto*, forms a well-defined homogenous taxonomic group even when analysed at the higher resolution of nucleotide composition. This is to be expected as crAssphage *sensu stricto* was the starting point for finding all crAss-like phages examined in this study and thus has the most sequences available for analysis.

**Figure 3.**
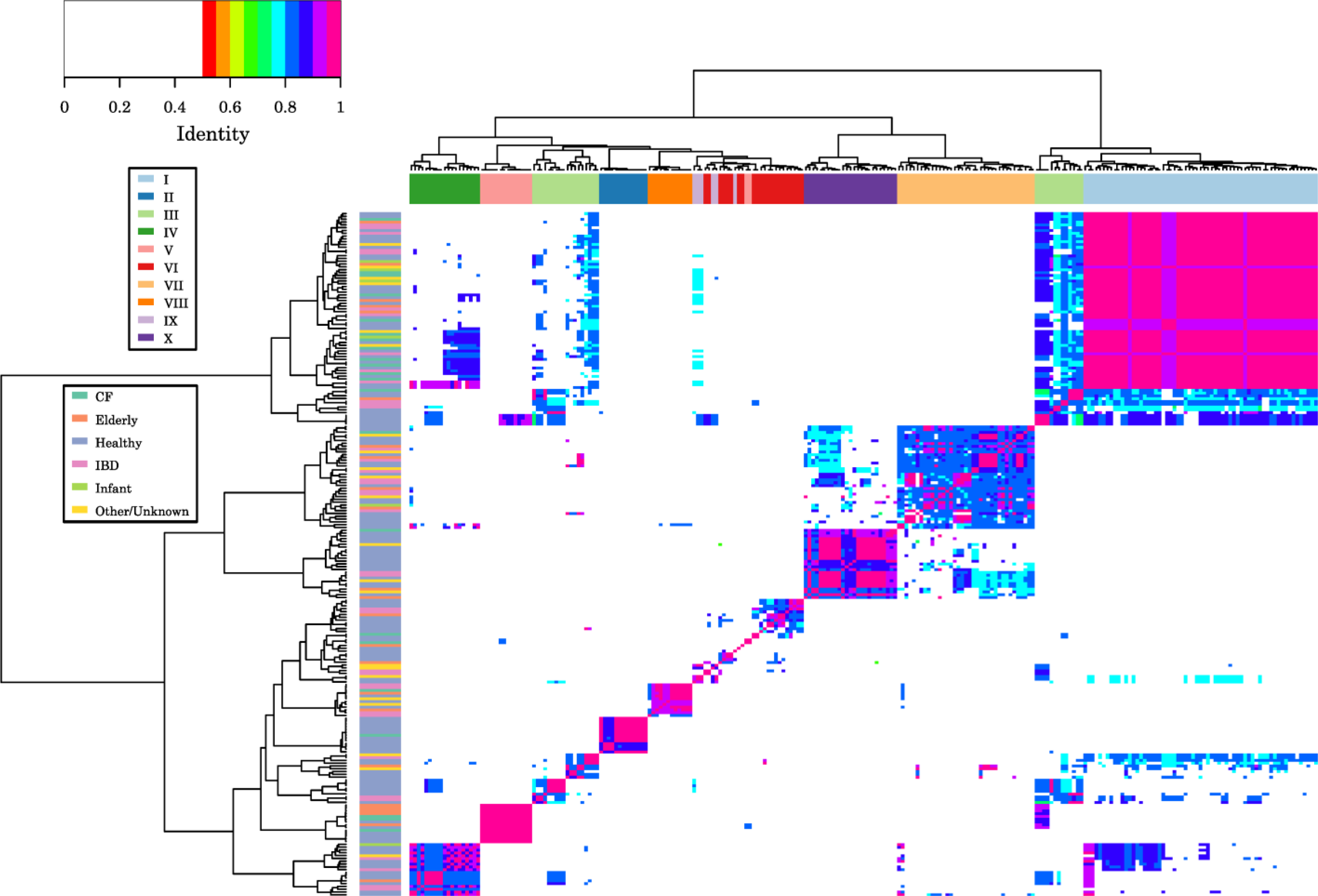
Average nucleotide identity of crAss-like phage contigs. The column annotation colour scheme highlights the predicted crAss-like phage candidate genus annotations, while the coloured row annotation represents the origin of the respective crAss-like phage contig.

Interestingly, the majority of crAss-like candidate genera demonstrate the same type of genomic organization (Supplementary Figure 2). Prominent features were shared between candidate genera I – V, IX, and X. These include; circular genomes with size ranging from 92 to 104kb, two clearly separated genome regions with opposite gene orientation and inversed G+C skew (the smaller region encodes proteins involved in replication, the bigger region coding for proteins involved in transcription and virion assembly, as suggested by Yutin *et al*.), the presence of giant open reading frames with sizes up to 15kb (UGP_052, UGP_053, UGP_052 in the genome of crAssphage *sensu stricto*), possibly coding for fused subunits of RNA polymerase (16), as well as an absence or scarcity of tRNA genes. By contrast, members of candidate genus VI had two genome regions of approximately equal size with opposite gene orientation and G+C skew and large sets of tRNA genes (up to 27; Supplementary Table 2). A prominent common feature of the members of candidate genera VII and VIII was absence of the giant open reading frames.

In order to further demonstrate the homogeneity of the candidate genus I of crAss-like phages, comparative genomic analysis was performed on complete genomes. We characterised crAss-like phages as having pac-type circularly permuted genomes (24, 25); therefore, only genomes determined as circular were considered for this analysis. The genomic start coordinates of circular Candidate Genus I crAss-like phages were altered to match that of the published prototypical crAssphage *sensu stricto*. Candidate Genus I crAss-like phages showed high levels of synteny and strong homology across their entire genomes. However, the most notable area of diversity is observed in the crAss-like phage putative receptor binding protein (UGP_074), which likely targets the different crAssphage strains towards their specific bacterial hosts (Supplementary Figure 3).

### Prevalence of crAss-like phages in human faecal virome samples

To get insights into relative abundance of different crAss-like phages in various human populations we aligned quality filtered reads, representing 532 human faecal samples from the same datasets as used for assembly of crAss-like genomes, to a database of 93 nonredundant crAss-like phage genomic sequences (with <90% of homology and/or <90% overlap between them) representing all 10 candidate genera.

Crass-like phage colonization rates varied from 51-58% in Malawian infants to 98-100% of healthy individuals of various ages in the Western cohorts. While relative crAss-like phage content ranged from 0 to 87% of the reads per sample, and depended significantly on the country of residence (p = 6.5E-09 in Kruskal-Wallis test) and age group of the donor (p = 1.6E-10). In ~8% of all virome samples, >50% of reads aligned to crAss-like phage genomes. Lowest overall crAss-like phage counts were seen in healthy Irish and Malawian infants and in USA adults with IBD (Figure 4A). On a global scale, crAss-like candidate genera I, III, and VIII seem to be the most prevalent ones (Figure 4B).

**Figure 4.**
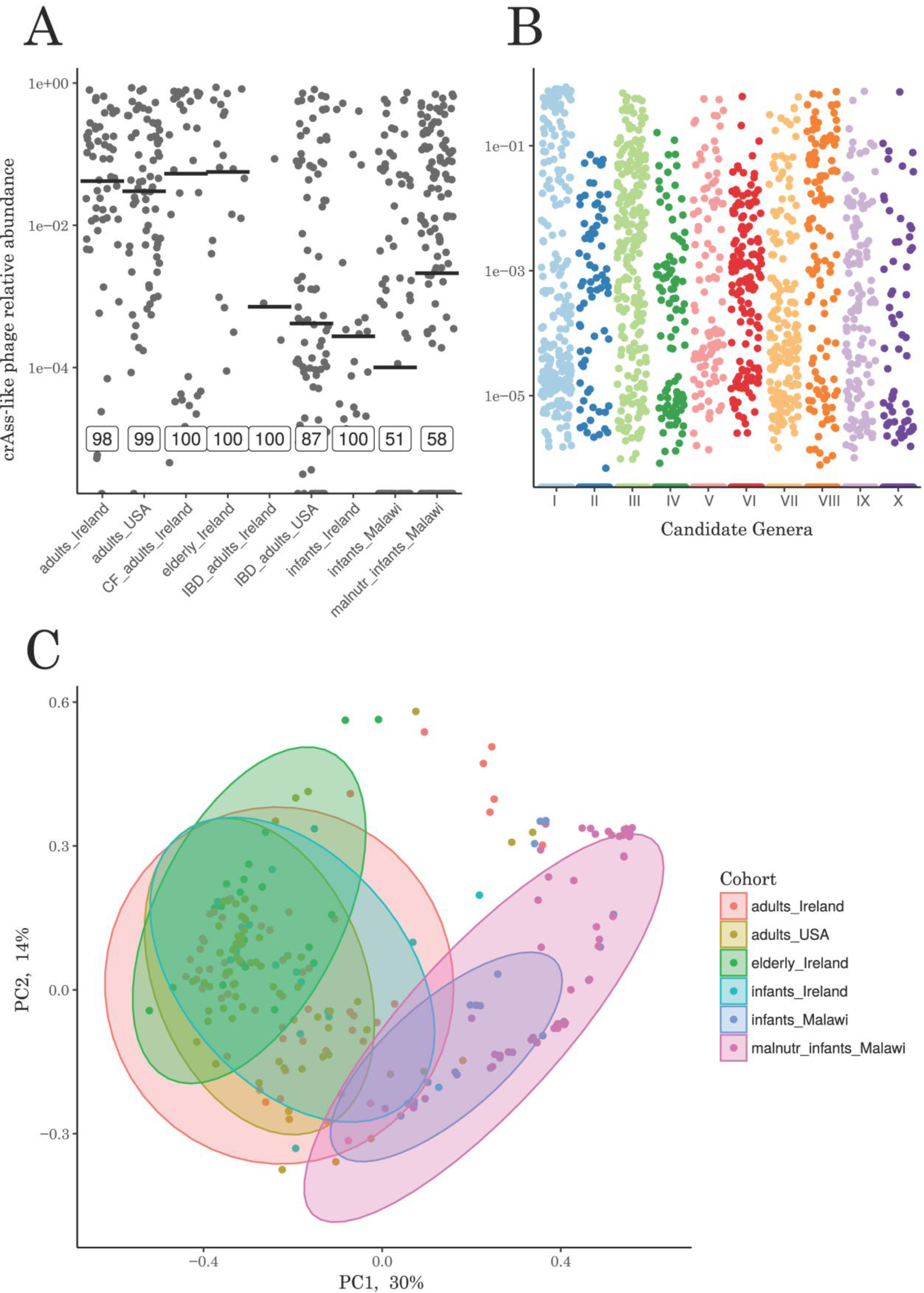
Prevalence of crAss-like phage in human faecal viromes. **(A)** Relative abundance of total crAss-like phage in several cohorts differing in age, health status and country of origin, based on the fraction of metagenomic reads aligned. Bars represent median relative abundances, the values within boxes represent percentage of positive samples. **(B)** Relative abundance of specific crAss-like candidate genera in total human populations analysed. **(C)** PCoA plot of crAss-like phages based on Spearman rank distances.

The specific composition of crAss-like phages in faeces partly separated a cohort of healthy and malnourished infants living in rural areas of Malawi from the healthy and diseased urban Western cohorts (Figure 4C). PERMANOVA analysis suggested that crAss-like phage composition was mostly driven by place of residence (R^2^ = 0.24, p = 0.001) with condition and age group also having significant impact (R^2^ = 0.05 and 0.01 respectively, p = 0.001). This observation is further supported by a clear difference in the distribution of specific crAss-like candidate genera across different populations (Figure 5). Specifically, Candidate Genus I, which includes crAssphage *sensu stricto* is by far the most prevalent type of crAss-like phages in Western population regardless of age. At the same time, same genus was extremely scarce in Malawian cohort where Candidate Genus III and VIII were the most common (p = 6.7E-03 and 1.4E-06, respectively).

**Figure 5.**
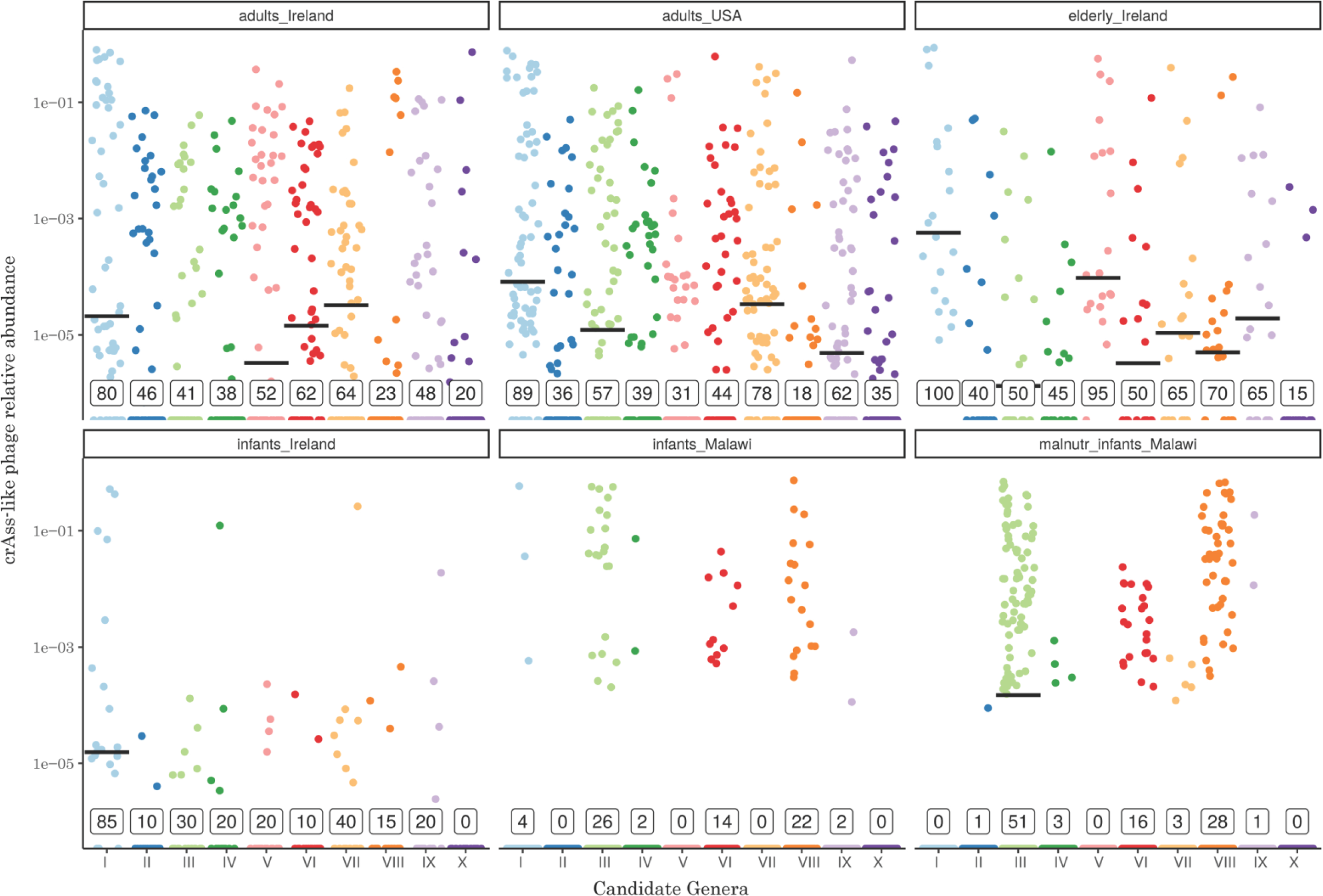
Relative abundance of the ten candidate genera of crAss-like phages in six different human cohorts based on the fraction of metagenomic reads aligned. Bars represent median relative abundances, while values within boxes represent percentage of positive samples.

### Faecal fermentations of a crAssphage rich sample

During an ongoing longitudinal study of faecal viromes in healthy adults we identified one individual (subject ID 924), in which crAssphage *sensu stricto* was consistently contributing >30% of virome metagenomic reads over a 12 month period. Thus, this donor was selected in order to investigate if crAssphage *sensu stricto* could be propagated in a batch faecal fermentation system. Quantitative PCR (qPCR) detection of a conserved fragment of the crAssphage *sensu stricto* DNA polymerase gene in the viral nucleic acid fractions throughout the fermentation revealed that crAssphage *sensu stricto* was effectively propagated. CrAssphage *sensu stricto* was found to increase in titre by 89 fold for up to 21 hours into the fermentation (Figure 6A).

**Figure 6.**
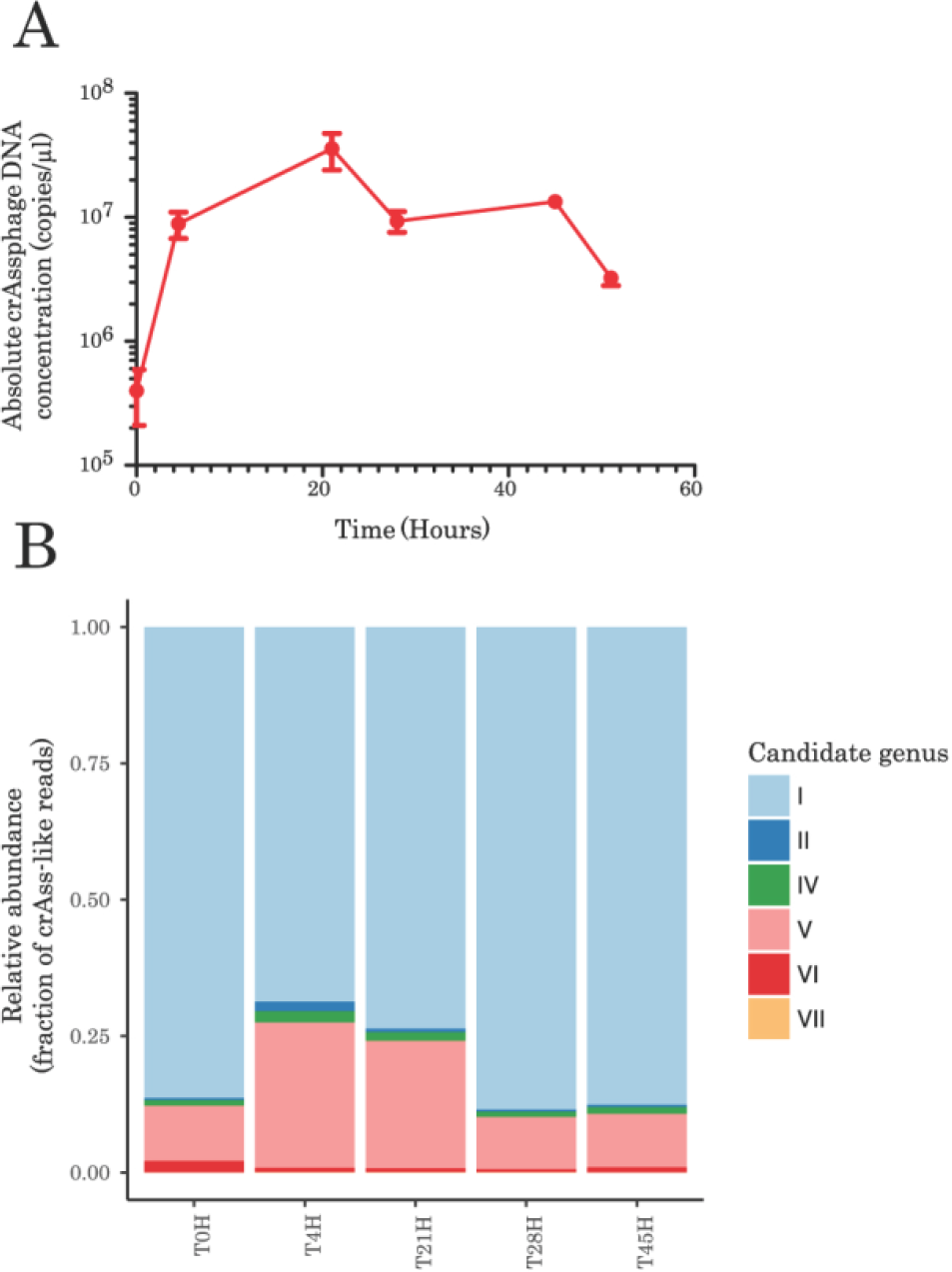
Analysis of crAss-like phage dynamics in a faecal fermenter. **(A)** Evidence of crAssphage *sensu stricto* propagation following *in vitro* fermentations (standard error, n=3). The level of crAssphage *sensu stricto* propagation was determined by qPCR analysis of viral-enriched DNA, respectively, using primers specific to a segment of the crAssphage *sensu stricto* DNA polymerase gene. **(B)** Six additional crAss-like phages, that group into five of the candidate genera, were identified following sequencing of the same viral-enriched DNA from the fermenter. The relative abundance of each of these crAss-like phages is skewed due to the biased amplification of other components of the viral-enriched DNA fraction that is associated with multiple displacement amplification.

Interestingly, shotgun metagenomic sequencing of the viral enriched DNA from the fermentation supernatants showed the presence of six other crAss-like phages in the study subject, in addition to crAssphage *sensu stricto* (Supplementary Table 2). These crAss-like phage contigs were all ≥70kb and grouped into five of the candidate genera (Figure 6B), four of which contributed to ≥1% of the reads per sample. The most abundant crAss-like contig of subject ID 924, designated as Fferm_ms_6 (linear, 90.4kb), is a member of proposed Candidate Genus I and closely related to crAssphage *sensu stricto*. Contig Fferm_ms_2 (linear, 88.8 kb) is the second most abundant in the sample and belongs to Candidate Genus V. Other crAss-like phages showed varying degrees of similarity at the amino acid level to different crAss-like phage at the genus-level taxonomic groups. Analysis of bacterial microbiota in the fermentation vessel using compositional 16S rRNA gene amplicon sequencing revealed a concomitant increase in the course of fermentation of a number of *Bacteroides* species, including; *B. dorei, B. uniformis, B. fragilis, B. xylanisolvens, B. nordii, Parabacteroides distasonis* and *Parabacteroides chinchillae* (Supplementary Figure 4).

### Biological characterisation of crAss-like phages

Transmission electron microscopy (TEM) of a crAssphage *sensu stricto* rich faecal filtrate showed a significant presence of short-tailed or non-tailed viral particles with icosahedral or isometric heads (53% of *Podoviridae* type and 29% of *Microviridae* or a smaller type of *Podoviridae*), with lower levels of tailed bacteriophages of the family *Siphoviridae* (15%; Figure 7A). *Podoviridae-*type virions could be further classified into two types: type I, with head diameters of ~76.5 nm and short tails; and type II, with a similar head size but head-tail collar structures and slightly longer tails (Figure 7B). Sequencing of the same fraction as used for the TEM showed that approximately 40% of reads aligned to crAss-like genomic contigs (Figure 7D). Based on the size of crAss-like genomic contigs assembled from subject ID 924 samples (88.8-97.3 kb), it seems likely that the predominant *Podoviridae* morphology observed corresponds to the crAss-like group of bacteriophages. For comparison, *Microviridae* phages have genomes 4.4-6.1 kb and icosahedral capsids of approx. 15-30 nm in diameter (26, 27).

**Figure 7.**
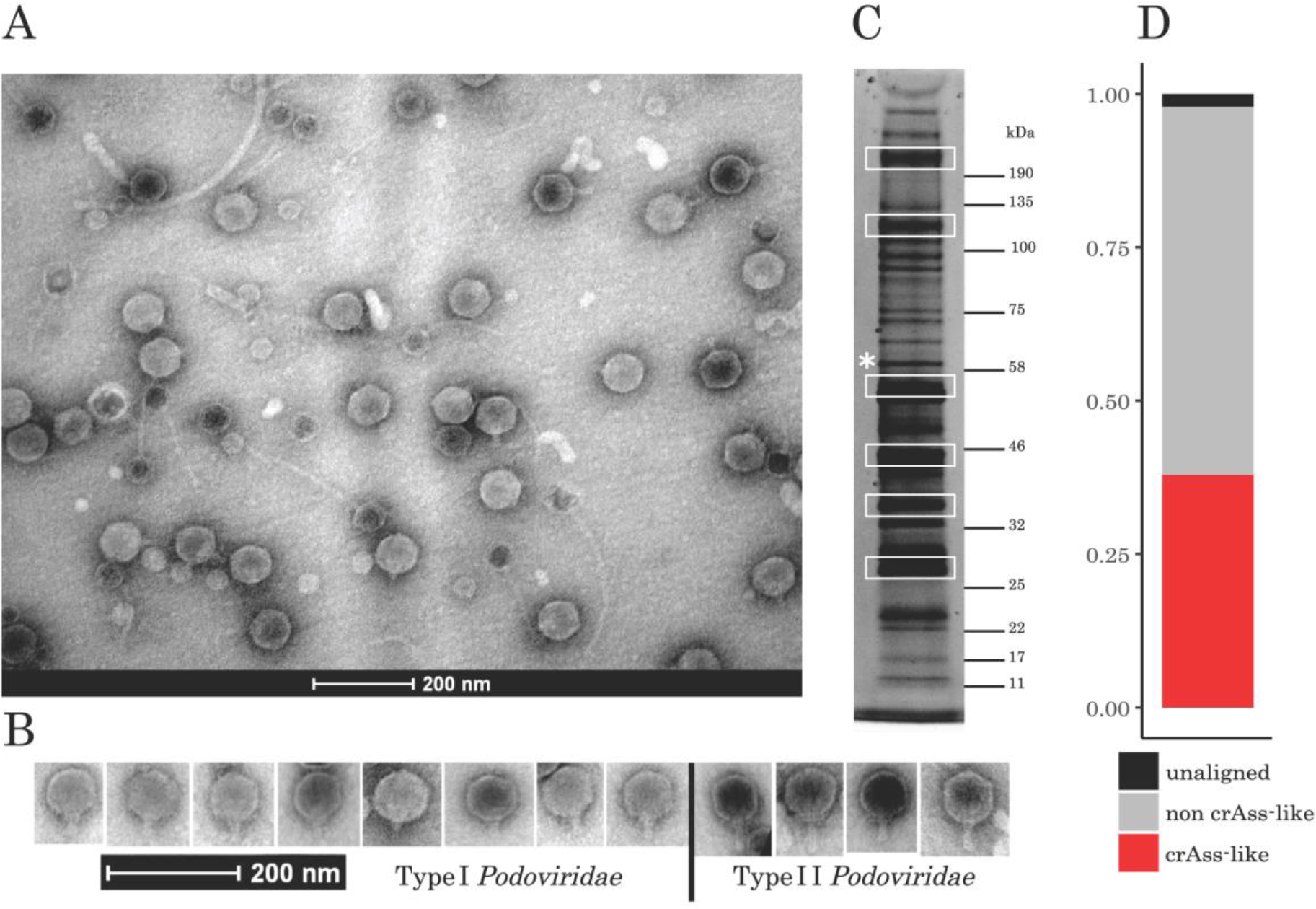
CrAss-like phage morphology was examined using a CsCl fraction purified from a crAssphage rich faecal filtrate of donor subject ID 924. **(A)** Analysis of the fraction through transmission electron microscopy (TEM) was performed. The TEM images are largely dominated by *Podovirdae* (53%), *Microviridae* (29%), *Siphoviridae* (15%) and other phage morphologies (3%). **(B)** Further examination of the observed *Podoviridae* identifies two variants with differing tail morphologies. Both variants have head diameters of ~76.5 nm. **(C)** SDS-PAGE gel of the CsCl fraction. Six bands containing possible crAssphage proteins were excised and analysed by mass spectrometry. A protein, denoted as Fferm_ms_2_MCP, isolated from the ~55 kDa (*) band was found to have high sequence similarity with Candidate Genus V crAss-like phages. **(D)** Sequencing of the CsCl purified viral fraction, without multiple displacement amplification, showed that approximately 40% the reads aligned to crAss-like phages.

The same CsCl fraction that was subjected to metagenomic sequencing and TEM visualisation was also analysed by SDS-PAGE followed by identification of major bands using MALDI-TOF mass spectrometry. A major structural protein of a crAss-like phage, denoted as Fferm_ms_2_MCP, was detected following MALDI-TOF analysis of a band excised from the ~55kDa area on a SDS-PAGE gel (Figure 7C). The obtained peptide profile corresponded to a protein of 490 amino acids and 55.4 kDa, encoded by Fferm_ms_2. Further analyses using BLASTp showed the protein to have 37% identity with UGP_086, predicted as the major capsid protein of the prototypical crAssphage (16).

In addition, we attempted to independently establish the size of crAss-like phage virions by passing faecal filtrates through a series of filters with gradually decreasing pore sizes (Supplementary Figure 5). Filtration through 0.1 μm pores (equivalent to 100 nm) resulted in partial retention of crAss-like phages while pores of 0.02 μm size completely removed crAssphage from the filtrate, as judged by the qPCR assay.

## Discussion

The overall objective of this study was to gain a more in depth insight into one of the most enigmatic phages discovered to date, crAssphage. This phage is highly abundant in the human microbiome on a global scale; however, it remains poorly understood. One reason why crAssphage has remained such a mystery is due to the lack of available genome sequences for comparison. When crAssphage was assigned a specific nomenclature and uploaded to a public repository by Dutilh and colleagues (12), it became a template for other studies to compare against. This highlights the need for researchers to upload both the sequencing reads and assembled contigs following metagenomic studies.

CrAssphage is a representative of an expanding group of human gut-associated bacteriophages. While previous studies have proposed a sequence-based classification of crAss-like viruses at the familial level (16), our *in silico* analysis fits within classical familial taxonomic assignments whereby crAss-like phages are categorised as *Podoviridae*. In this study, we present 243 new crAss-like phage genomes from various metagenomic studies. Comparative genomics of the 244 available crAss-like phages demonstrates an extensive degree of diversity among these phages, including the potential identification of four crAss-like phage subfamilies. While the *alphacrAssvirinae* subfamily is currently the largest of the 4 subfamilies, future studies looking for additional homologues of *betacrAssvirinae*, *gammacrAssvirinae* and *deltacrAssvirinae* members will refine these taxonomic categories.

Assigning phage taxonomy, in the absence of a universal genetic marker such as 16S rRNA, is a difficult and potentially erroneous process. In our study, we adopted a method previously employed to assign taxonomic ranks to *Podoviridae* based on the percentage of shared homologous proteins (17). This categorisation strategy identified 10 candidate genera, with crAss-like phages in each genera originating from the faeces of putatively healthy individuals and people suffering from various diet and bowel-related disorders. Alternative proposed methods for defining phage genera include grouping phages with >50% nucleotide similarity identity together (18). Noteworthy, the 10 proposed crAss-like phage genera as determined by percentage of shared homologous proteins closely resembles that observed for crAss-like phage groups when characterised by >50% shared average nucleotide identity.

Several crAss-like phage genera proposed in this study have distinct nucleotide G+C compositions. The nucleotide composition of obligate parasites, such as phages, likely evolves in close association with the host bacterium (23, 28–30). Thus, Candidate Genera III and VI with diverse G+C compositions are either heterogeneous groups of crAss-like phages that require further sequences to refine their taxonomic structure, or they are potentially capable of infecting across a broad host range.

Quantitative analysis of crAss-like phage content in several cohorts revealed that in agreement with the previous studies the vast majority of faecal viral metagenomic samples contained varied amounts of crAssphage DNA. CrAssphage *sensu stricto* (Candidate Genus I) is by far most predominant type in Western populations, co-existing with other crAss-like phages in the majority of samples. By contrast, in the cohort of malnourished and healthy Malawian infants (9, 31), other candidate genera such as III, VIII and IX seem to play the leading role. It is well known that non-Western rural populations, which mostly consume high fibre, low fat and low animal protein diet are predominantly associated with high *Prevotella/*low *Bacteroides* type of gut microbiota (known as enterotype II (32)), as opposed to *Bacteroides*/*Clostridia*-dominated microbiota (enterotype I) in urban populations consuming western diet (33, 34). Indeed, our analysis of the Reyes et al. (2015) 16S rRNA gene sequencing data confirmed high prevalence of *Prevotella* in Malawian samples (Supplementary Figure 6). One can hypothesize that members of candidate genera III, VIII and IX might be associated with *Prevotella* or other members of the order *Bacteroidales* apart from *Bacteroides sensu stricto*.

The *in vitro* analysis of samples obtained from subject ID 924 was particularly intriguing. By mapping metagenomic sequencing reads against crAssphage *sensu stricto*, it was initially thought that this donor only carried the prototypical crAssphage at levels exceeding 30% of total viral reads for a 1 year period. A subsequent mining for phages related to crAssphage *sensu stricto* using metagenomic sequencing at later time points, with and without multiple displacement amplification resulted in 5 additional crAss-like phages being simultaneously detected from a single donor. However, the initial screening and inclusion criteria for bioinformatic detection of crAss-like phages resulted in a fragmented crAss-like phage contig being missed. The overlooked crAss-like phage, Fferm_ms_2 (Candidate Genus V), turned out to be extremely important during the *in vitro* biological characterisation experiment. Therefore, it is possible many additional crAss-like phage genomes could be present within the metagenomic datasets that were examined in this study, but they were not included in our analysis because of the inclusion criteria chosen or even the choice of assembly program.

In total, subject ID 924 consistently carried 7 crAss-like phages, which resolved in our taxonomic analysis into 5 candidate genera. Three of the crAss-like phages were identified in Candidate Genus VI, supporting the notion this is a heterogeneous group and not simply composed of broad host range infecting phages. It is possible that there are potentially more than 7 crAss-like phages within subject ID 924. However, we believe that only a single representative of each candidate crAss-like phage genus (with the exception of the heterogeneous candidate genus VI) could assemble correctly, with two or more highly identical phages amalgamating their single nucleotide polymorphisms into a single consensus representative sequence (Supplementary Figure 7).

This study demonstrates the proliferation of crAss-like phages in a faecal fermenter, the first evidence of crAss-like phage propagation in the laboratory. Furthermore, following our ability to propagate faecal crAss-like phages, we conducted the first transmission electron micrographs (TEMs) of these phages. Indeed, the most abundant faecal viruses present in samples used to inoculate faecal fermentation were *Podoviridae*. This is in agreement with the predictions made by Yutin *et al*., following their detailed genome annotation of two crAss-like phages (16). Interestingly, however, our TEMs suggest presence of two types of virions with short non-contractile tails (Figure 7C). Presumably, the more abundant type I virions with shorter tail can belong to members of Candidate Genus I, also found as the most abundant crAss-like phage group in subject ID 924 by means of metagenomic sequencing (Figure 6B). Whereas type II virions with slightly longer tails and visible head-tail collar structures may correspond to Candidate Genus VI, found as the second most abundant crAss-like phage subfamily in shotgun metagenomics. But without isolating these phages in pure culture, it is not possible to accurately assign which *Podoviridae* tail corresponds to which specific crAss-like phage subfamily or genera.

This work provides the first *in vitro* evidence confirming that crAss-like phages are members of the *Podoviridae* family. This is shown from three levels of experimentation using the same CsCl fraction purified from crAssphage rich faeces of a healthy human donor. The TEM images produced from the CsCl fraction showed an abundance of the signature *Podoviridae* morphology. Other phage capsids present, predominantly *Microviridae*, would typically be associated with smaller genome sizes than that of crAss-like phages (26). Sequencing of the same fraction identified that almost 40% of the reads aligned to crAss-like phages. This is consistent with the percentage of *Podoviridae* identified in the TEM images. Furthermore, a highly predominant protein denoted as Fferm_ms_2_MCP, was isolated from the fraction and was found to have significant similarity to crAss-like phages of (Candidate Genera V) as well as a moderate degree of similarity to crAssphage *sensu stricto* (Candidate Genera I). This *in vitro e*vidence, in line with the taxonomic analysis performed by Yutin *et al*., proves that crAss-like phages do indeed belong to the *Podoviridae* family.

Identifying a means of propagating crAss-like phages is of particular importance. However, it was also observed that the primers applied in the qPCR analyses of viral nucleic acids were not suitable for targeting crAss-like phages associated with the various subfamilies and candidate genera that differed significantly from crAssphage *sensu stricto*. With the availability of more crAss-like phage sequences, broad and narrow spectrum primers can now be designed and applied in the analysis of these phages. The choice of primers for detecting crAss-like phages was also discussed in the recent work of Cinek *et al*. (14). This will be an important part of further work.

It also has to be considered that human gut crAssphage is not one single entity, but rather a group of diverse viruses, sharing certain signature genomic traits. It is most likely that these diverse phages target multiple bacterial taxa. Previously, a member of the *Bacteroides* genus was hypothesised as being the host for crAssphage (12). In a study prior to the discovery of crAssphage (35), a 95.9kb contig corresponding to a putative virus φHSC05 was shown to be stably engrafted after transplantation of human faecal virus fraction into germ-free mice colonized with an artificial defined community of 15 bacterial species. The artificial bacterial community, among others, included: *Bacteroides thetaiotaomicron* (2 strains), *B. caccae*, *B. ovatus*, *B. vulgatus*, *B. cellulosilyticus* and *B. uniformis*. One might conclude that one of the above mentioned 7 strains of the genus *Bacteroides*, more likely than the remaining 8 strains of Gram-positive anaerobic bacteria used in that study, must have served as a host for crAssphage propagation. The retrospective analysis of contigs from that study conducted by ourselves showed that the φHSC05 contig was 91.73% identical by its nucleotide sequence to crAssphage *sensu stricto*. Since crAssphage had not been described at the time the article was published, this very interesting observation was never made by the authors of the original work.

With more divergent sequences, we could assume that different members of the *Bacteroides* genus, or even *Bacteroidetes* phylum for example, may serve as hosts for different crAss-like phages. One host that has been hypothesised for prototypical crAss-like phages is *B. dorei*. This was inferred following the analysis of a dataset generated from infants and toddlers with islet autoimmunity. It was correlated that crAssphage was only present when *B. dorei* also was detected within the samples. This was not true for other *Bacteroides* members tested, including *B. vulgatus* which is highly related to *B. dorei*. This correlation is compelling; however, it should be noted that there was no confirmation that crAssphage has any role in causing bacteriome alterations that lead to islet autoimmunity (36). Interestingly, one of the key *Bacteroides* species detected from our faecal fermentation 16S rRNA analysis was *B. dorei*. Its levels were inversely proportional to that of crAssphage. Therefore, this possible phage-host pair should be investigated further.

CrAss-like phages have also been defined as a part of the core human gut phageome (10). This emphasises the importance of identifying hosts for diverse crAss-like phages belonging to different candidate genera proposed in this study. Such knowledge along with the ability to propagate crAss-like phages *in vitro* will provide an insight into its biological significance including their possible role in shaping the bacterial composition of the human gut microbiome in a positive or negative manner, in context of various disease states, such as inflammatory bowel disease, cancer, and obesity among others. Thus far, only a few studies has attempted to correlate crAss-like phages with a gastrointestinal disorder (7, 13, 36). Exploring this aspect of crAss-like phages further will be a key part of future work.

In conclusion, our results expand the repertoire of known crAss-like phages significantly, providing a path towards the identification of further crass-like phages and their hosts. This will lead to a better understanding of their role, if any, in human health and disease. Our work also provides an interesting insight into the diversity of these human gut-associated phages in various populations through *in silico* and *in vitro* methods. In addition, we also demonstrate that these enigmatic phages can be efficiently propagated *in vitro* in a mixed culture as well as present the first TEMs of crAss-like phages, giving an insight into their morphology. CrAss-like phages appear to be universally present in human populations, including various disease states. Due to the specificity of phage-host interactions, the diversity of crAss-like phages suggests they infect multiple diverse bacteria of the human gastrointestinal microbiota. However, more studies will be required to determine the biological significance and role of crAss-like phages in the human gut and determine if its presence positively or negatively impacts human gastrointestinal health.

## Methods

### Metagenomic datasets and contig assemblies

Sequencing reads from publicly available metagenomic datasets were downloaded from NCBI Sequence Read Archive (SRA) database. All published and unpublished metagenomic datasets that yielded crAss-like phage contigs, the DNA preparation protocol, the sequencing technology, the assembly program, and information related to contig nomenclature, are briefly described in Supplementary Table 1. All reads were processed using Trimmomatic v0.32 to remove adaptor sequences and to trim reads when the Phred quality score dropped below 30 for a 4bp sliding window. Trimmed reads were assembled using either SPAdes v3.6.2 (37) or metaSPAdes v3.10.0 (38). Contigs from the assembly of 702 metagenomic samples were assigned a specific nomenclature, representing: [1] study/sample description, [2] SPAdes or metaSPAdes assembly, and [3] numerical rank of largest-to-smallest assembled contigs. The full list of contigs assembled in this study, the available associated metadata, and contig accession numbers, are detailed in Supplementary Table 2.

### Detection and curation of crAss-like phages

The detection of crAss-like phage contigs was performed as follows. The amino acid polymerase sequence of prototypical crAssphage (UGP_018, NC_024711.1) was queried using BLAST v2.2.28+ (39) against a translated nucleotide database consisting of assembled metagenome contig sequences. The most conserved orthologous protein group detected in our initial putative crAss-like phage screening included prototypical crAssphage protein UGP_092, which was annotated through the HHPred homology and structural prediction web server (40) as a phage terminase. This was then used as a second genetic signature of crAss-like phages and used in an additional BLAST search. All putative crAss-like phages selected for analysis met the following criteria: [1] a BLAST hit against either prototypical crAssphage polymerase or terminase with an e-value less than 1e-05, [2] a BLAST query alignment length ≥350bp, and [3] a minimum contig length of 70kb (representing near-complete crAss-like phage contigs).

### Identification of crAss-like phage orthologous proteins and clusters

The encoded proteins of crAss-like phages were predicted using Prodigal v2.6.3 (41). Orthologous proteins shared between crAss-like phages were detected using OrthoMCL v2.0 using default parameters (42). The presence/absence of orthologous proteins between crass-like phages was initially converted into a binary count matrix where the percentage of shared orthologous proteins was calculated (Figure 1B). The optimum number of phage clusters was calculated using the percentage of shared homologous proteins using the NbClust v3.0 package for R (43). Hierarchical clustering was performed on the count matrix of percentage shared crAss-like phage orthologous proteins using Ward’s minimum variance method [‘Ward.D2’ algorithm in R (44)]. The resulting dendrogram was cut at k = 10 based on the estimation of the number of crAss-like phage clusters (Figure 1A).

As a verification of the 10 predicted crAss-like phage clusters, the original abundance matrix of crass-like phage orthologous proteins was used to calculate Euclidean distances between samples. These distance variations were calculated using the t-SNE machine learning algorithm [‘tsne’ v0.1-3 for R; (45)] and plotted using ggplot v2.2.1 (Figure 2). The presence or absence of orthologous protein groups was used to determine the core proteome of crAss-like phage clusters (Supplementary Figure 8).

### Phylogeny of crAss-like phage terminase sequences

Following the work of Yutin *et al*., (16) all publically available crAss-like phage terminase sequences were included in an additional phylogenetic analysis (Supplementary Figure 2). The terminase amino acid sequences of crAss-like phages were aligned using Muscle v3.8.31 (46). The resultant alignment was converted to Phylip format and phylogeny was determined by PhyML using a JTT amino acid substitution model (47). The phylogenetic tree was visualised using FigTree v1.4.3. The phylogenetic tree is coloured based on the crAss-like phage clustering analysis with node support values displayed.

### Genomic comparisons of crAss-like phages

The average nucleotide identity between crAss-like phage contigs was calculated using Pyani v0.2.3 by the ANIm method with a 500bp fragment size. Pairwise comparisons of complete crAss-like phage genomes belonging to Candidate Genera I was performed using Easyfig v2.2.2. Genomic start coordinates and contig orientations were altered to match the published GenBank sequence of prototypical crAssphage NC_024711.1. The order of crAss-like phages in the Easyfig image was adjusted to match to the order they appear in the average nucleotide identity analysis (Figure 3). The Easyfig image was generated using tBLASTx comparisons, with a minimum BLAST length of 50bp and identity of 30bp (Supplementary Figure 3). The presence of crAss-like phage tRNA-encoding sequences were detected using ARAGORN v1.2.36 (48). To determine the genomic packaging mechanism of crAss-like phages, metagenomic sequencing reads from a TruSeq (Illumina) manually fragmented DNA library were analysed using PhageTerm (25). Single nucleotide polymorphisms (SNPs) of crAss-like phages were observed by aligning metagenomic sequencing reads to the consensus assembled contig sequence using Bowtie2 and Samtools, and visualising SNPs using Tablet v1.17.08.17 (49).

### Alignment of virome metagenomic reads to crAss-like contigs

The quality filtered reads from 532 human faecal viromes (as subset of 701 viromes selected based on availability of sufficient metadata) were then aligned to the set of 93 nonredundant crAss-like phage genomic (with <90% of homology and/or <90% overlap between them) using Bowtie2 v2.3.0 (50) using the end-to-end alignment mode. A count table was generated with Samtools v0.1.19 which was then imported into R v3.3.1 for statistical analysis. β-diversity of crAss-like viral populations in human cohorts was visualized using PCoA plot based on Spearman rank distances (D = 1 – ρ, where ρ is Spearman rank correlation coefficient of relative abundance of different crAss-like contigs between samples). Statistical analysis was performed using permutational multivariate analysis of variance (PERMANOVA) implemented in Vegan v2.4.3 package for R (51) and non-parametric Kruskal-Wallis test.

### Recruitment of a crAssphage faecal donor and faecal fermentations

Human faecal viromes from a number of ongoing studies sequenced using Illumina HiSeq and MiSeq platforms were screened for crAss-like phages by aligning the obtained sequencing reads against prototypical crAssphage NC_024711.1 using Bowtie2 v2.3.0. One individual (subject ID 924) was found to carry crAssphage consistently at levels exceeding 30% of the total number of reads over a one year period. The recruited individual is an adult female that suffers from gastritis and is vitamin B12 deficient. A frozen standard inoculum (FSI) sample was processed as described by (52) with the following modification: the sample was resuspended in 1X phosphate buffered saline (37 mM NaCl, 2.7 mM KCl, 8 mM Na_2_HPO_4_, and 2 mM KH_2_PO_4_.), 0.05% (w/v) L-cysteine (Sigma Aldrich, Ireland) and (1 mg/L) resazurin (Sigma Aldrich, Ireland). The crAssphage-rich FSI was inoculated into 400 ml YCFA-GSCM broth in a 500 ml fermenter vessel at 5% (v/v). Fermentation media was prepared exactly as described by (53) with the addition of glucose (2 g/L), soluble starch (2 g/L), cellobiose (2 g/L) and maltose (2 g/L). Fermentation was performed in batch format at approximately 37°C for 51 hours. Dissolved oxygen was sustained at <0.1% by constantly sparging the vessel with anaerobic gas mix (80% (v/v) N_2_, 10% (v/v) CO_2_, 10% (v/v) H_2_) and stirring at 200 rpm. Both 2M NaOH and HCl solutions were used to maintain pH at ~7. Samples were collected at the following time points; 0, 4, 21, 28, 45 and 51 hours. Collected samples were centrifuged at 4,700 rpm at +4°C for 10 minutes. The resulting supernatants were filtered once through a 0.45 μM pore syringe filter and stored at +4°C. Resultant pellets were stored at -80°C.

### Extraction of viral nucleic acids and sequencing library preparation

Total virome extractions were performed on 0.45 μM pore filtered fermentation supernatants. Solid NaCl and polyethylene glycol 8000 were added to the filtrates to give a final concentration of 0.5M and 10% (w/v), respectively. After overnight incubation at +4°C samples were centrifuged at 4,700 rpm and +4°C for 20 minutes. The pellets were then resuspended in 400μl of SM buffer (1M Tris-HCl pH 7.5, 5M NaCl, 1M MgSO_4_) and briefly vortexed with an equal volume of chloroform. This mixture was then centrifuged at 2,500g for 5 minutes using a standard desktop centrifuge. The resultant aqueous phase was then transferred into an Eppendorf to which 40μl DNase buffer (10mM CaCl_2_ and 50mM MgCl_2_) and 8U and 4U TURBO DNase (Ambion/ThermoFisher Scientfic) and RNase I (ThermoFisher Scientific) were added, respectively. This was incubated at 37°C for 1 hour followed by an enzyme inactivation step at 70°C for 10 minutes. This was followed by the addition of 2μl proteinase K and 10% SDS and further incubation at 56°C for 20 minutes. Lastly, 100μl phage lysis buffer (4.5 M guanidinium isothiocyanate, 44 mM sodium citrate pH 7.0, 0.88% sarkosyl, 0.72% 2-mercaptoethanol) was added to lyse the viral particles. The final incubation was carried out at 65°C for 10 minutes. The resulting lysates were lightly vortexed with an equal volume of phenol/chloroform/isoamyl alcohol 25:24:1 (Fisher Scientific) and were centrifuged at room temperature for 5 minutes at 8,000g. This was again repeated with the resulting aqueous phase. Following the second extraction, the aqueous phase was passed through a DNeasy Blood and Tissue Kit (Qiagen) for final lysate purification. The wash steps were each repeated twice and the final elution was carried out in 50μl elution buffer. Viral DNA quantification was carried out with the Qubit HS DNA Assay Kit (Invitrogen/ThermoFisher Scientific) in a Qubit 3.0 Flurometer (Life Technologies). The viral nucleic acids were then subjected to reverse transcription using SuperScript IV Reverse Transcriptase (RT) kit (Invitrogen/ThermoFisher Scientific). The protocol was carried out exactly as described in the manufacturer’s protocol for random hexamer primers. Following this, 1μl of the reversed transcribed viral DNA was subjected to GenomiPhi V2 (GE Healthcare) Multiple Displacement Amplification (MDA). Finally, MDA and non-MDA viral DNA was prepared for sequencing using TruSeq DNA Library Preparation Kit (Illumina, Ireland). All steps were performed as per the manufacturer’s instructions. Prepared libraries were sequenced on an Illumina HiSeq platform (Illumina, San Diego, California) with 2x300bp paired-end chemistry at GATC Biotech AG, Germany. Reads were filtered, trimmed and assembled into contigs as described above. A count matrix was created by aligning quality-filtered reads back to contigs using Bowtie2 and Samtools.

### CrAssphage PCR detection

Two oligonucleotide primer pairs were designed based on the prototypical crAssphage DNA polymerase sequence UGP_018 (1) using PerlPrimer software (54). Primer sequences are as follows: CrAss-Pol-F5 5’-GCCTATTGTTGCTCAAGCTATTGAA-3’, CrAss-Pol-R5 5’-ACAACAGAACCAGCTGCCAT-3’, CrAss-Pol-F6 5’-AGTGGTCTTGCTCCNGAACAATGG-3’ and CrAss-Pol-R6 5’-AACCTCCAGTTGCAACAGTATAAGT-3’. PCR products were cloned into pCR2.1-TOPO TA vector (ThermoFisher Scientific) and obtained plasmids at known concentrations were used to establish calibration curves through serial two-fold dilutions. Subsequently, qPCR were run in 15μl reaction volumes using SensiFAST SYBR No-ROX mastermix and LightCycler 480 thermocycler with the following conditions: initial denaturation at 95°C for 5 minutes, then 35 cycles of 94°C for 20 seconds, 55°C for 20 seconds and 72°C for 20 seconds, with a final extension at 72°C for 7 minutes. All samples were run in triplicate and the standard error was determined following calculation of DNA concentration based on the above standard curve.

### Electron microscopy and detection of crAssphage proteins

A virus-enriched fraction of the crAssphage positive faecal sample, collected from subject ID 924, was prepared for electron microscopy imaging as follows. A 1:20 suspension (w/v) of faeces was prepared in SM buffer followed by vigorous vortexing until homogenised. The homogenised sample was chilled on ice for 5 minutes prior to centrifugation twice at 4,700 rpm for 10 minutes at +4°C. The resulting supernatant was then filtered twice through a 0.45 μM pore syringe filters. The filtrate was ultra-centrifuged at 120,000g for 3 hours using a F65L-6x13.5 rotor (ThermoScientific). The resulting pellets were resuspended in 5 ml SM buffer. The viral suspensions were ultracentrifuged again by overlaying them onto a caesium chloride (CsCl) step gradient of 5M and 3M, followed by centrifugation at 105,000g for 2.5 hours. A band of viral particles visible under side illumination was collected and buffer-exchanged using 3 sequential rounds of 10-fold diluting and concentrating to the original volume by ultra-filtration using Amicon Centifugal Filter Units 10,000 MWCO (Merck). The purified fraction was then analysed by qPCR for the presence of crAssphage as described above. Following this, 5μl aliquots of the viral fraction were applied to Formvar/Carbon 200 Mesh, Cu grids (Electron Microscopy Sciences) with subsequent removal of excess sample by blotting. Grids were then negatively contrasted with 0.5% (w/v) uranyl acetate and examined at UCD Conway Imaging Core Facility (University College Dublin, Dublin, Ireland) by transmission electron microscope. The faecal viral fraction from subject ID 924 was further concentrated using Amicon Ultra-0.5 Centrifugal Filter Unit with 3 kDa MWCO membrane (Merck, Ireland). This concentrated fraction was loaded onto a premade Bolt 4-12% Bis-Tris Plus reducing SDS-PAGE gel (Invitrogen) and separated at 200 V for 30 minutes using 1X NuPAGE MOPS SDS Running Buffer. Six brightest bands with approximate molecular weights of 28, 35, 45, 55, 120 and 200 kDa were excised and subjected to MALDI-TOF/TOF (Bruker ultraflex III) protein identification following in-gel trypsinization, at Metabolomics & Proteomics Technology Facility (University of York, York, UK).

### 16S rRNA gene library preparations

Total DNA was extracted from the pellets formed following centrifugation of fermentation samples. This was carried out using the QIAamp Fast DNA Stool Mini Kit (Qiagen, Hilden, Germany). All steps were carried out as per the manufacturer’s protocol with the addition of a bead-beating step to aid total DNA extraction from the bacterial cells. Approximately 200mg of each pellet was placed in a 2ml screw-cap tube containing a mixture of one 3.5 mm glass bead, a 200μl scoop of 1mm zirconium beads and a 200μl scoop of 0.1mm zirconium beads (ThistleScientific) with 1ml of InhibitEX Buffer. Bead-beating was carried out three times for 30 seconds using the FastPrep-24 benchtop homogeniser (MP Biomedicals). Between each bead-beating the samples were cooled on ice for 30 seconds. The samples were then lysed at 95°C for 5 minutes. All other steps were carried out as per the manufacturer’s protocol. Following extraction of total bacterial DNA, the hypervariable regions of V3 and V4 16S ribosomal RNA genes were amplified from 15ng of the DNA using Phusion High-Fidelity PCR Master Mix (ThermoFisher Scientific) and 0.2µM of each of the following primers, containing Illumina-compatible overhang adapter sequences: 16S-FP: 5’-TCGTCGGCAGCGTCAGATGTGTATAAGAGACAGCCTACGGGNGGCWGCAG-3’ and 16S-RP: 5’-GTCTCGTGGGCTCGGAGATGTGTATAAGAGACAGGACTACHVGGGTATCTAATCC-3’. The PCR program was run as follows: 98°C for 30 seconds, 25 cycles of 98°C for 10 seconds, 55°C for 15 seconds and 72°C for 20 seconds, with a final extension of 72°C for 5 minutes. The amplicons were then purified using Agencourt AMPure XP magnetic beads (Beckman-Coulter) followed by a second PCR to attach dual Illumina Nextera indices using the Nextera XT index kit v2 (Illumina). Purification was performed once again and the libraries were quantified using a Qubit dsDNA HS Assay Kit. The libraries were then pooled in equimolar concentration and sent for sequencing on an Illumina MiSeq platform (Illumina, San Diego, California) at GATC Biotech AG, Germany. The quality of the raw reads were assessed with FastQC (v11.5) and initial quality filtering was performed using Trimmomatic v0.36. Filtered reads were imported into R (v3.4.3) for analysis with DADA2 v1.6.0. (55) Further quality filtering and trimming (maxN of 0 and a maxEE of 2) was carried out on both the forward and reverse reads with only retention in cases of pairs being of sufficient high quality. Error correction was performed on forward and reverse reads separately and following this, reads were merged. The resulting unique Ribosomal Variant Sequences (RSVs) were subjected to further chimera filtering using USEACH v8.1 (56) with the Chimera-Slayer gold database v20110519. The retained, high quality, chimera-free, RSVs were classified with the RDP-classifier in mothur v1.34.4 (57) against the RDP database v11.4 (phylum to genus) and SPINGO (58) for species assignment. Plots were generated using the R package ggplot2 v2.2.1.

## Acknowledgements

This publication has emanated from research conducted with the financial support of Science Foundation Ireland (SFI) under Grant Numbers SFI/12/RC/2273, SFI/15/ERCD/3189 and SFI/14/SP APC/B3032, and a research grant from Janssen Biotech, Inc.

## Author contributions

EG and SRS performed the laboratory and bioinformatic work, respectively. AS assisted in both the laboratory and bioinformatic analyses. AGC performed the 16S analysis. FJR, TDSS, LAD and EGT assisted in the design, implementation and interpretation of experiments. EG, AS and SRS wrote the paper and generated the figures. AGC, FJR, TDSS, LAD and EGT reviewed drafts of the manuscript and provided constructive criticism for its improvement. PR and CH secured the funding and wrote the paper. All authors contributed to the analysis of the data.

## Data deposition

The 244 crAss-like phage contigs analysed in this study have been submitted to GenBank and are currently under revision. Contigs are currently accessible at: https://figshare.com/articles/crAss-like_contigs_fasta_tar_gz/6098321

---

**Supplementary Figure 1.**
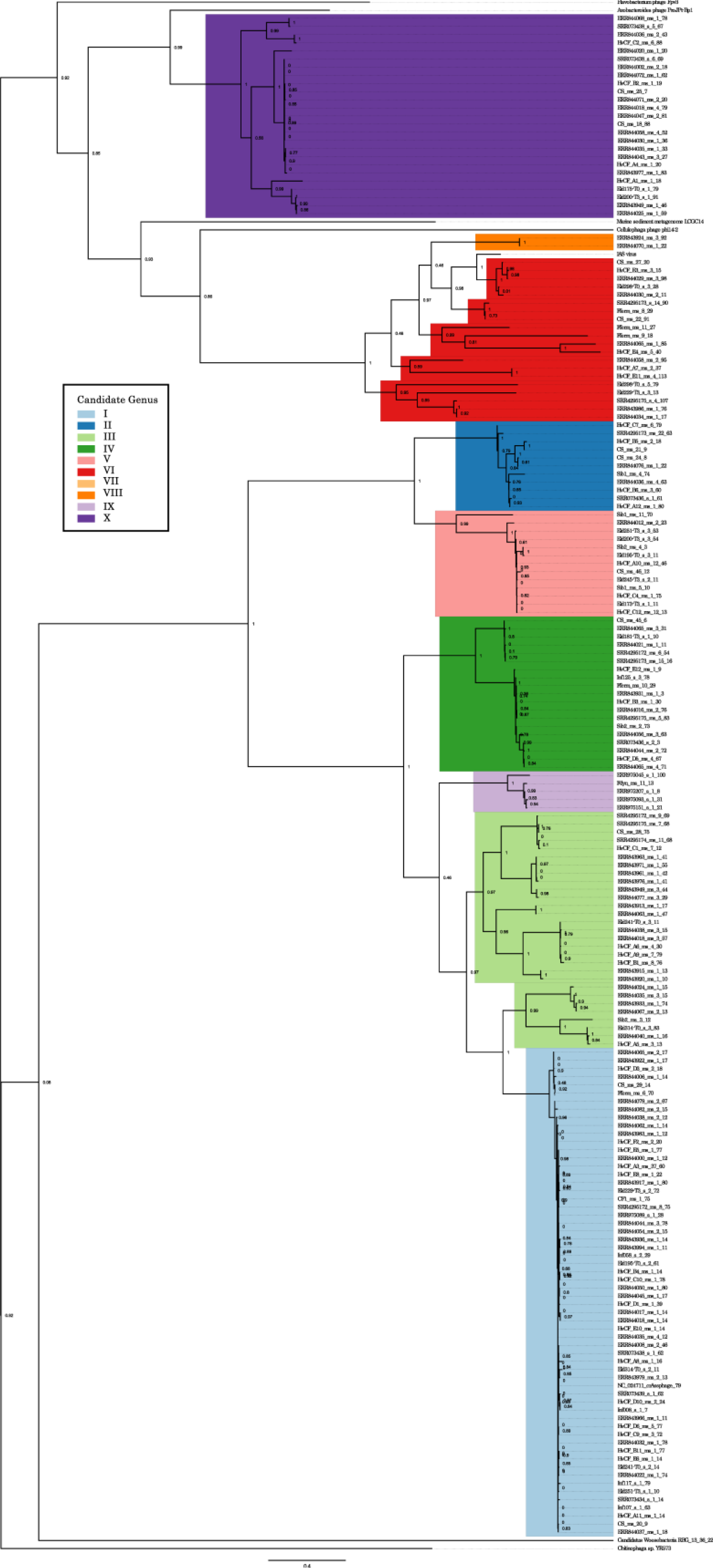
Phylogeny of crAss-like phage terminase protein sequences, including publically available terminase sequences from the Yutin *et al*. (2017) characterisation of familial-related crAss-like phages. The figure legend insert corresponds to the colour scheme of the 10 proposed candidate genera groupings. NC_024711 crAssphage and IAS virus, discussed in the main text, are highlighted in red. Bootstrapping node support values are shown.

**Supplementary Figure 2.**
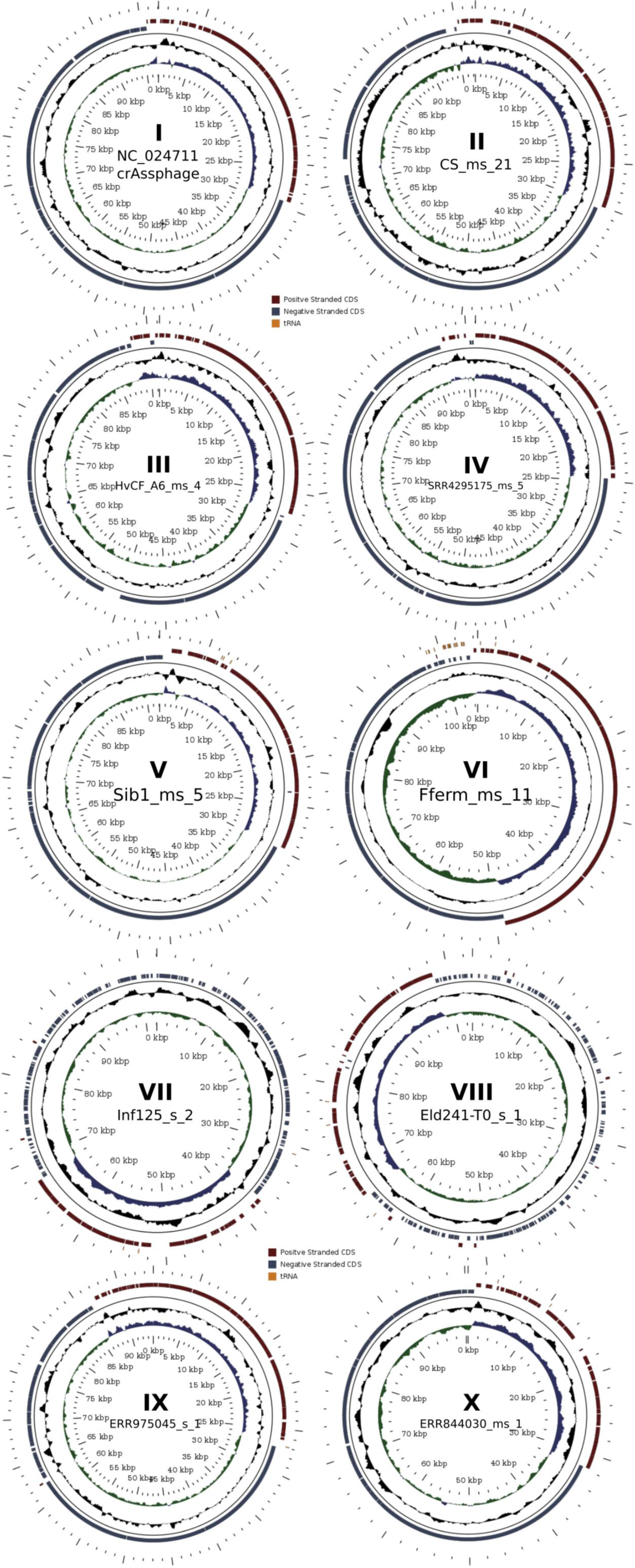
Comparison of general structural feature of representative complete circular genomes of the 10 proposed genera of crAss-like bacteriophages. Innermost circle (green/blue), G+C skew; middle circle, G+C content deviation from mean value; outermost circle, protein-coding genes (CDS) located on positive (red) and negative (blue) DNA strands, respectively; and tRNA genes (orange).

**Supplementary Figure 3.**
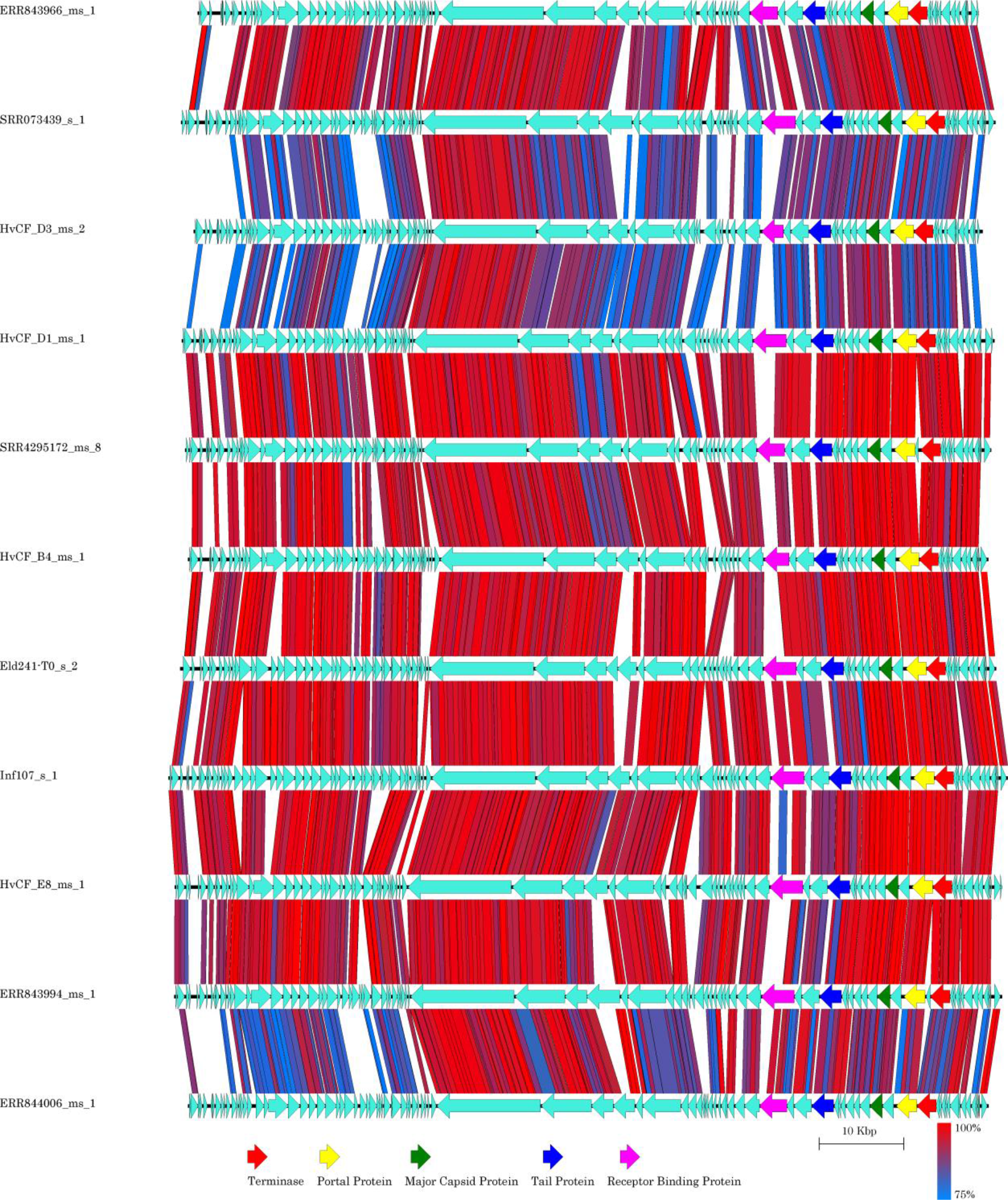
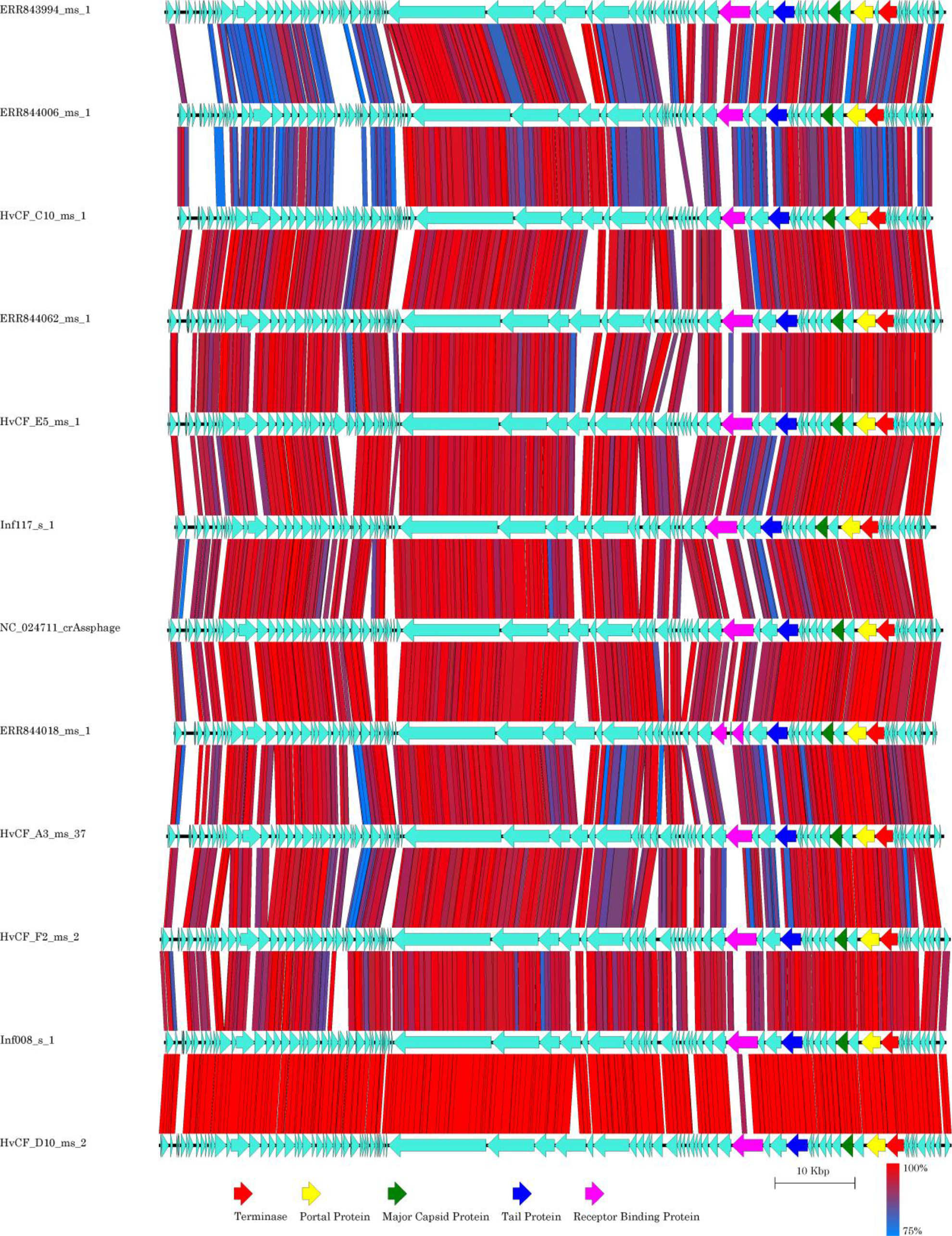
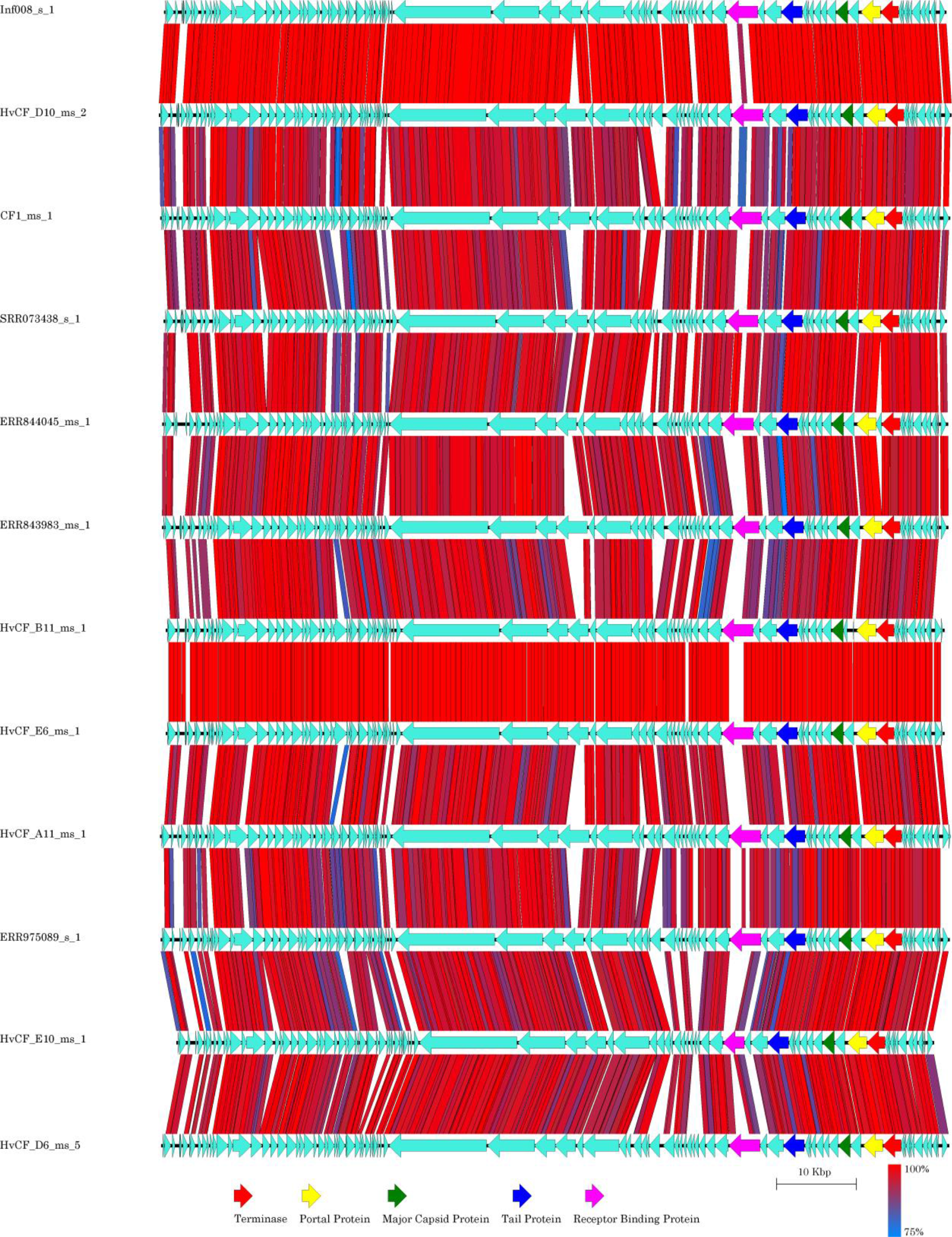
Comparison of circular Candidate Genus I crAss-like phage genomes. Start co-ordinates of crAss-like phage genomes were adjusted to match crAssphage *sensu stricto*. The order of crAss-like phage genomes was determined by the average nucleotide identity comparisons. Open reading frames corresponding to specific predicted phage structural proteins are highlighted.

**Supplementary Figure 4.**
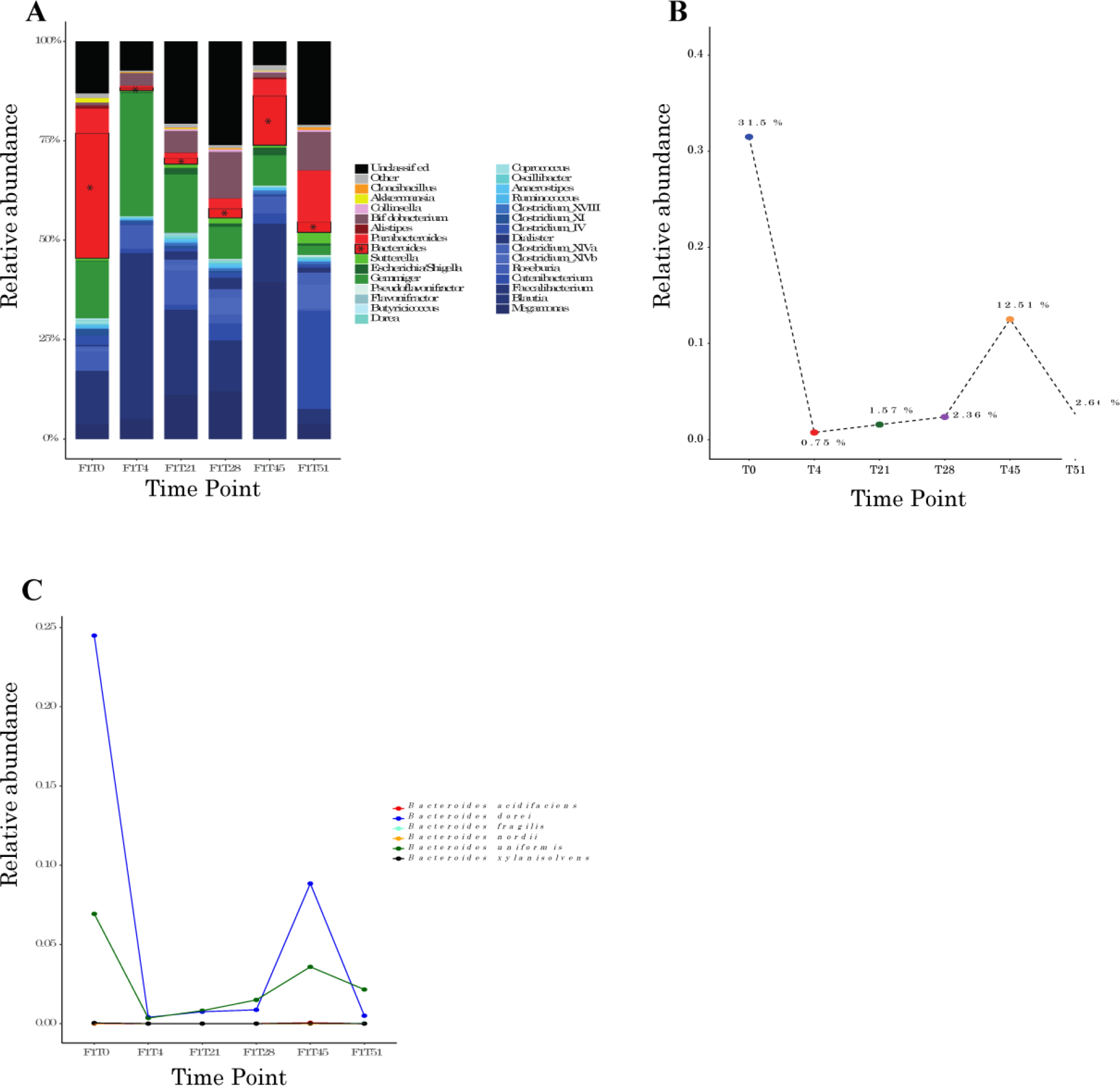
The relative abundance of 16S rRNA throughout the crAssphage-rich frozen standard inoculum initiated faecal fermentation. **(A)** The relative abundance of the major genera detected throughout the fermentation. *Bacteroides* (*), the genus hypothesised to be associated with crAssphage, can be seen to decrease between time points 0 and 4 of the fermentation after which levels gradually begin to increase again. **(B)** The relative abundance of total *Bacteroides* at each time point. **(C)** Abundances of individual *Bacteroides* species detected. *B. dorei* is found to be particularly abundant and seemingly inversely proportional to the detected crAssphage levels.

**Supplementary Figure 5.**
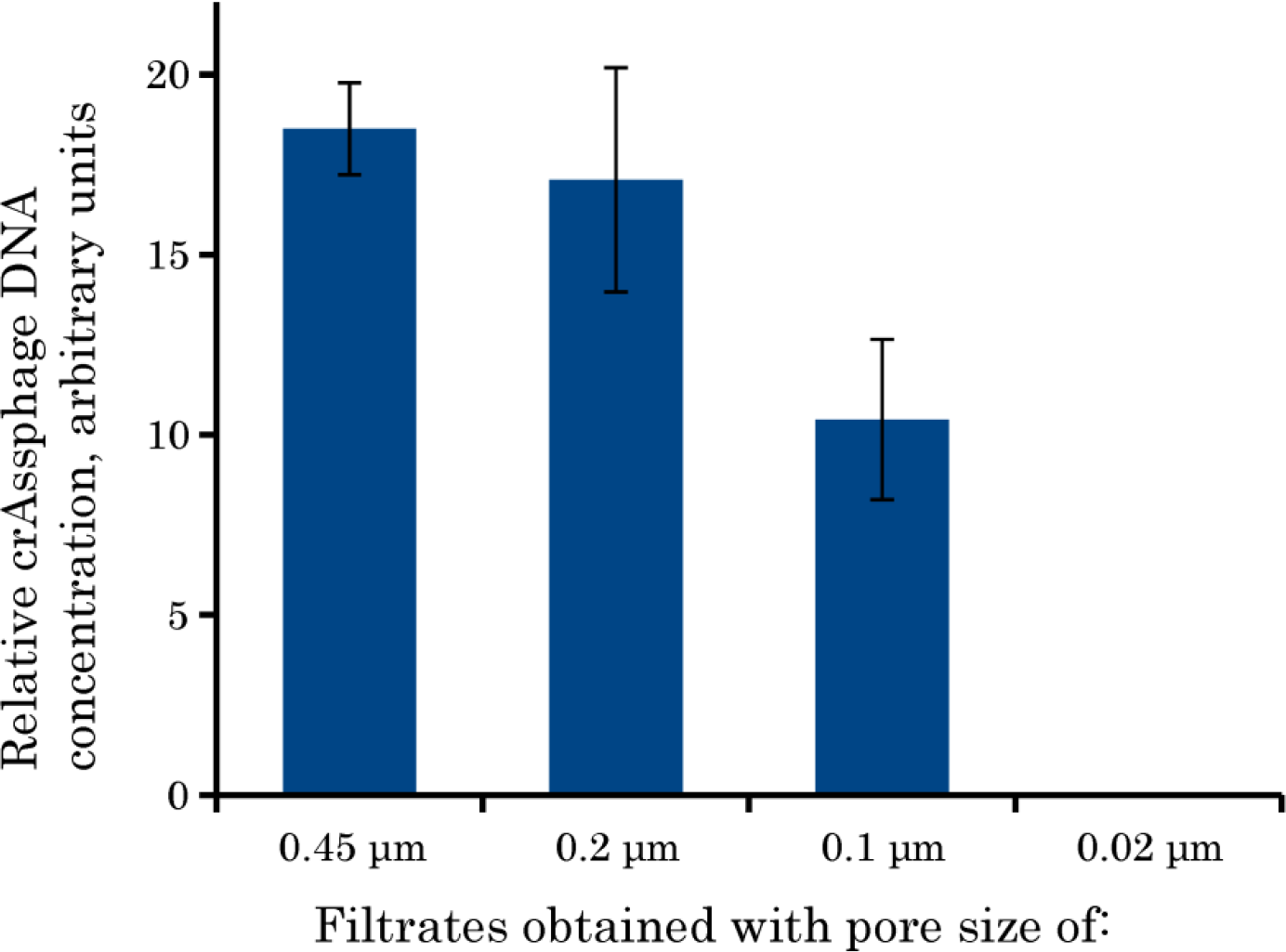
Quantitative PCR analysis of filtrates obtained with different pore sizes from a crAssphage-rich faecal sample collected from subject ID 924.

**Supplementary Figure 6.**
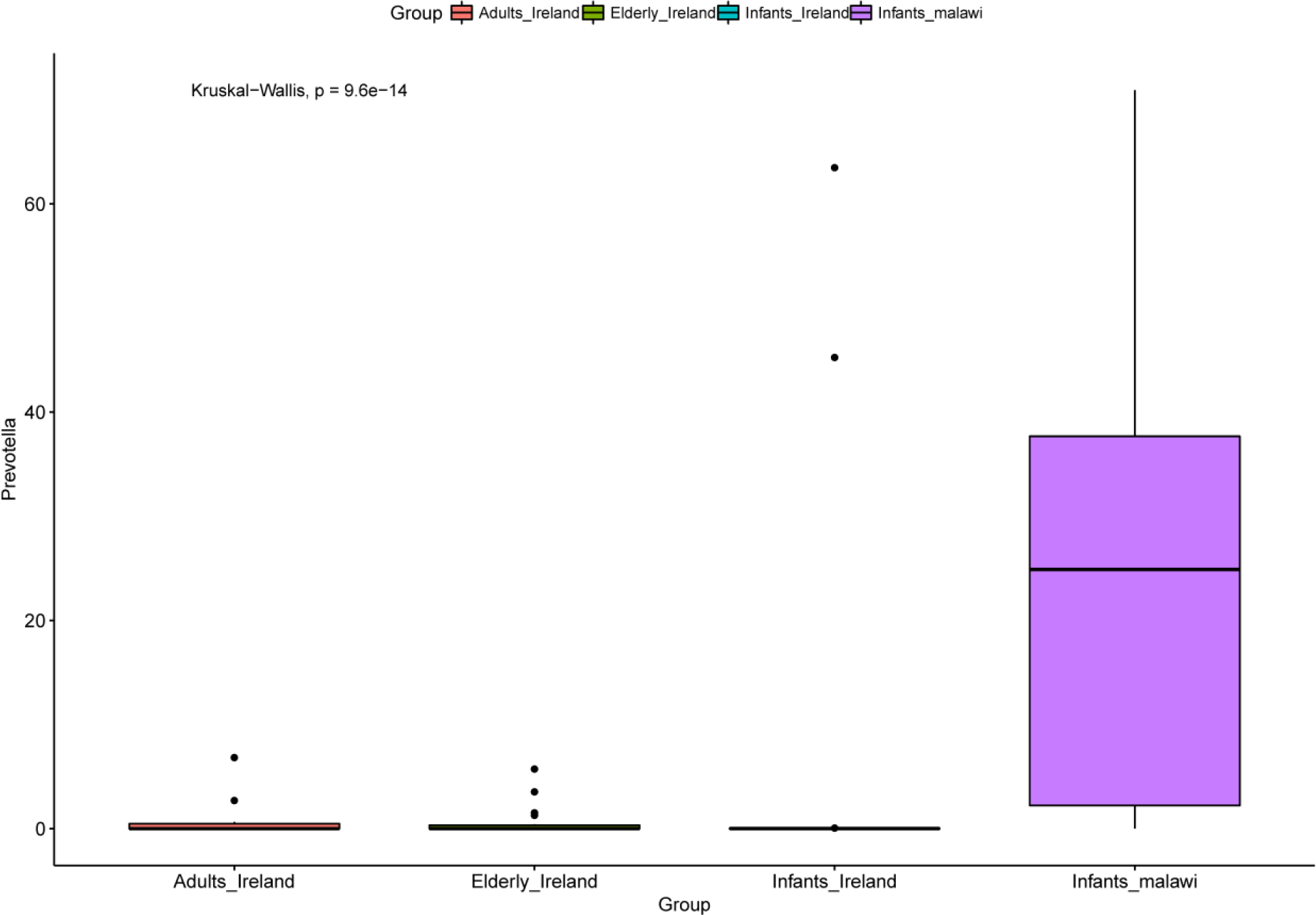
Comparison of 16S rRNA Prevotella abundances in healthy Irish adults and infants with Malawian infants.

**Supplementary Figure 7.**
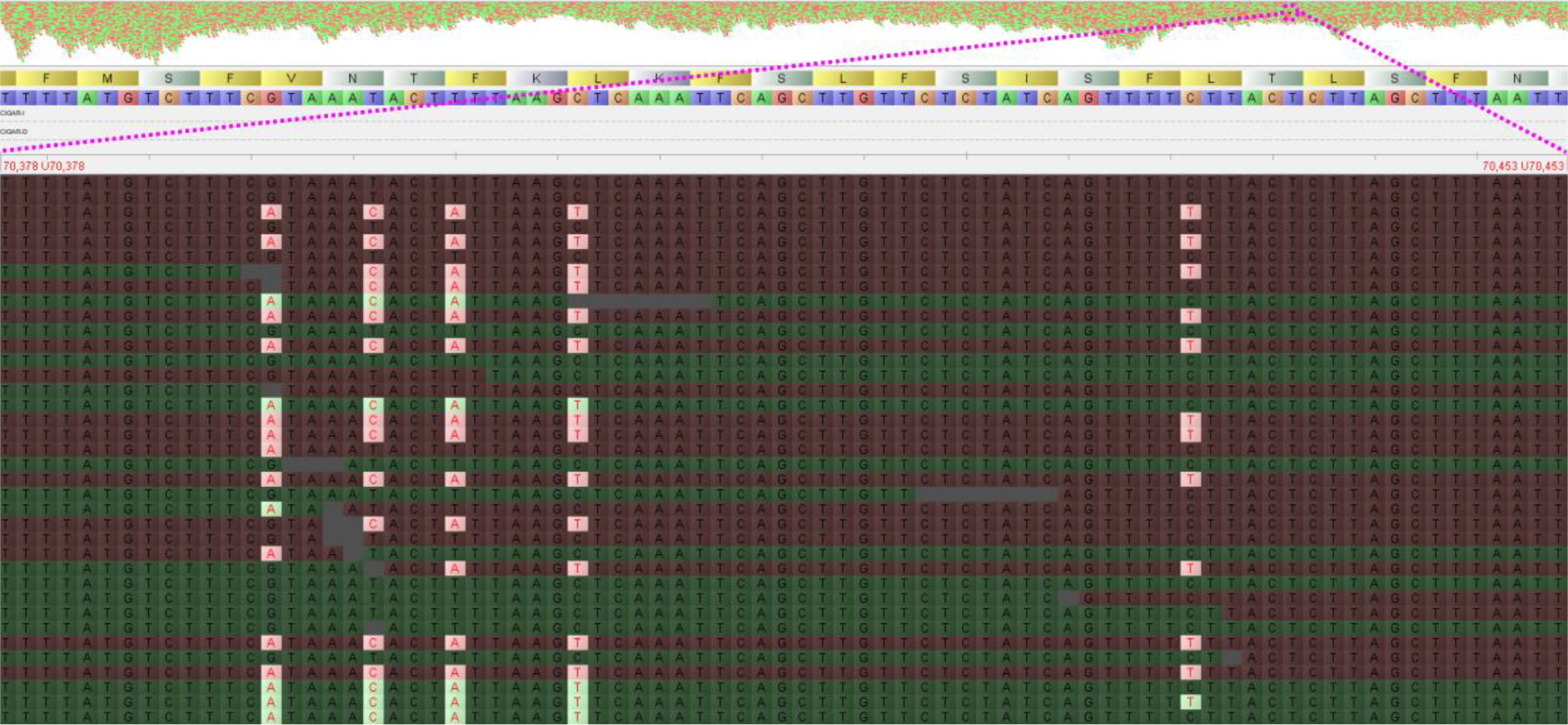
Visualisation of an example of metagenomic read-specific single nucleotide polymorphisms within the assembled of crAss-like phage contig, Fferm_ms_2, highlighting within sample species and/or strain level diversity of crAss-like phages are not resolved.

**Supplementary Figure 8.**
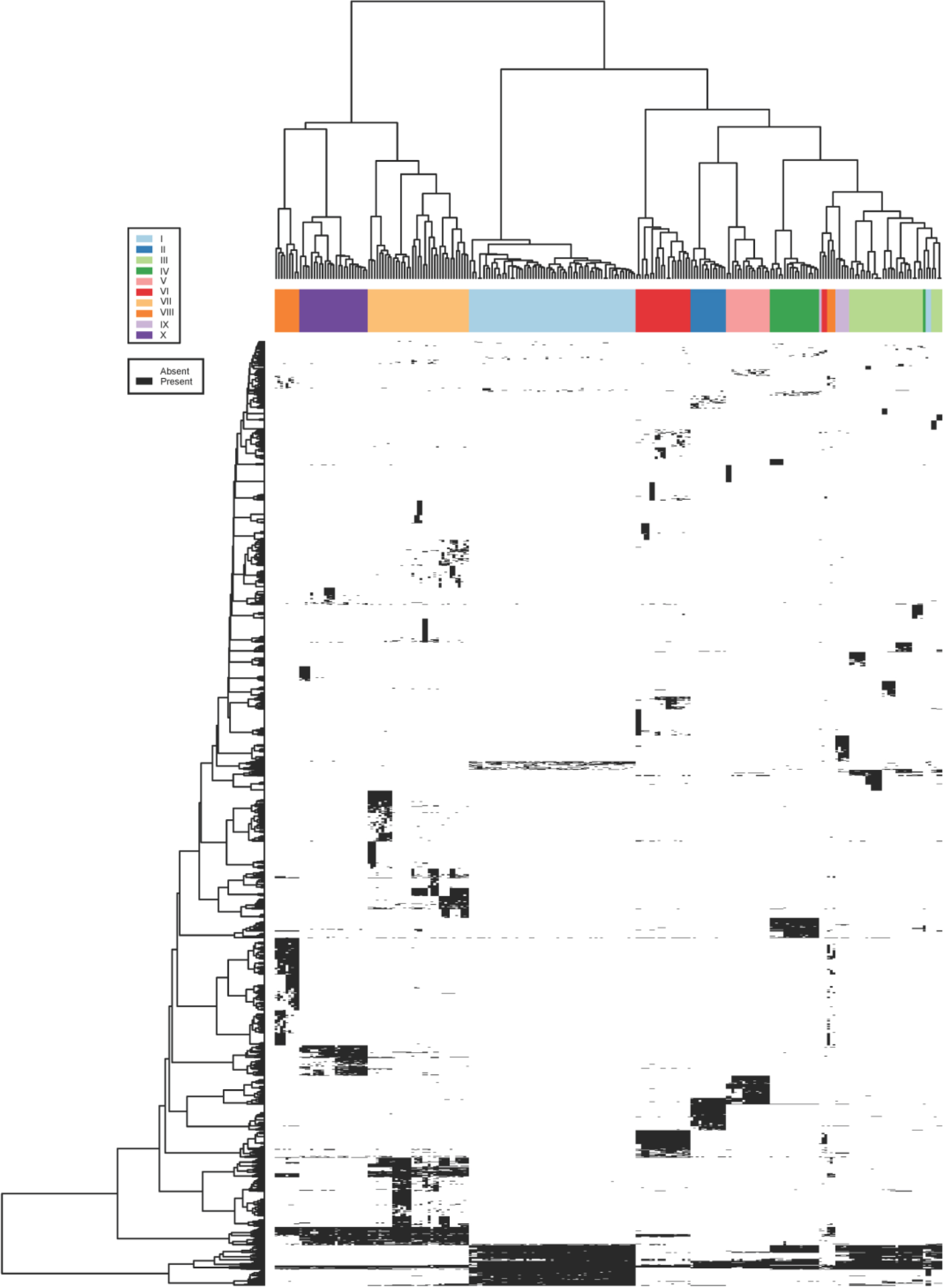
Visualisation of the core proteome of the 10 crAss-like phage candidate genera.

